# Dozer: Debiased personalized gene co-expression networks for population-scale scRNA-seq data

**DOI:** 10.1101/2023.04.25.538290

**Authors:** Shan Lu, Sündüz Keleş

## Abstract

Population-scale single cell RNA-seq (scRNA-seq) datasets create unique opportunities for quantifying expression variation across individuals at the gene co-expression network level. Estimation of co-expression networks is well-established for bulk RNA-seq; however, single-cell measurements pose novel challenges due to technical limitations and noise levels of this technology. Gene-gene correlation estimates from scRNA-seq tend to be severely biased towards zero for genes with low and sparse expression. Here, we present Dozer to debias gene-gene correlation estimates from scRNA-seq datasets and accurately quantify network level variation across individuals. Dozer corrects correlation estimates in the general Poisson measurement model and provides a metric to quantify genes measured with high noise. Computational experiments establish that Dozer estimates are robust to mean expression levels of the genes and the sequencing depths of the datasets. Compared to alternatives, Dozer results in fewer false positive edges in the co-expression networks, yields more accurate estimates of network centrality measures and modules, and improves the faithfulness of networks estimated from separate batches of the datasets. We showcase unique analyses enabled by Dozer in two population-scale scRNA-seq applications. Co-expression network-based centrality analysis of multiple differentiating human induced pluripotent stem cell (iPSC) lines yields biologically coherent gene groups that are associated with iPSC differentiation efficiency. Application with population-scale scRNA-seq of oligodendrocytes from postmortem human tissues of Alzheimer disease and controls uniquely reveals co-expression modules of innate immune response with markedly different co-expression levels between the diagnoses. Dozer represents an important advance in estimating personalized co-expression networks from scRNA-seq data.

## Introduction

The advent of single-cell RNA sequencing (scRNA-seq) has provided unparalleled insights into the transcriptional programs of cell types and cellular stages (Zeisel et al., 2015; Chen et al., 2017; Travaglini et al., 2020). Emerging population-scale scRNA-seq datasets are enabling investigations of population-level phenotypic variability as a function of transcriptomic variability at the single cell level (Bernardes et al., 2020; Jerber et al., 2021; van der Wijst et al., 2020; Cuomo et al., 2020; Soskic et al., 2022). When combined with individual-level genetic information, population-scale scRNA-seq datasets enable mapping expression quantitative trait loci (eQTL) across different cell types and in dynamic processes (Van Der Wijst et al., 2018; Soskic et al., 2022).

A key opportunity unveiled by emerging scRNA-seq datasets is the construction of personalized gene co-expression networks which can be leveraged to link network-level properties to phenotypic variation, e.g., discovering therapeutic targets in cancer (Forbes, 2022) and identifying genetic variants (e.g., network QTLs) that associate with network properties such as modules (Langfelder and Horvath, 2008) and network centrality (Savino et al., 2020) measures. Gene co-expression network analysis (Zhang and Horvath, 2005), which estimates gene-gene correlations, is a key inference tool for detecting latent relationships that might be obscured in standard analysis of clustering and differential expression. Research on protein-protein interactions and co-expression networks has established that genes with high centrality are crucial for survival and specific processes related to the studied phenotype (He and Zhang, 2006; Jeong et al., 2001; Lareau et al., 2015; Azuaje, 2014). Looking beyond high centrality genes in a single network, detecting genes with differential centrality between phenotypic groups can reveal regulatory alterations. Savino et al. (2020) discussed that genes with differential centrality, but no differential expression, may indicate potential regulatory relationships. Such differences in gene centrality may arise due to modifications in post-translational processes, involvement of a cofactor, or epigenetic mechanisms, such as DNA methylation, which can modify the regulatory function of a transcription factor (TF) without impacting its expression. Workflows, including data preprocessing, normalization, and network transformation, for estimation of gene-gene correlations in co-expression networks are well-established for bulk RNA-seq (Johnson and Krishnan, 2022); however, single-cell measurements of expression pose unique challenges due to technical limitations and noise levels inherent to the technology. Past research innovated numerous approaches to mitigate the noise and sparsity issues related to scRNA-seq technology when estimating gene-gene correlations. MetaCell (Baran et al., 2019) aggregated profiles of disjoint and homogeneous groups of cells to reduce sparsity in expression counts. bigSCale2 (Iacono et al., 2019) leveraged patterns of differential expression between clusters of cells to calculate correlations from transformed scRNA-seq data. State-of-the-art scRNA-seq data normalization and imputation methods, such as SCTransform (Hafemeister and Satija, 2019), MAGIC (Van Dijk et al., 2018), SAVER (Huang et al., 2018) and DCA (Eraslan et al., 2019) perform well in estimating expression, removing technical variability, and improve downstream dimension reduction and clustering tasks. However, these widely used methods, notably except for SAVER, rarely account for the estimation uncertainty in their expression estimates and were observed to introduce correlation artifacts for gene pairs that are not expected to be co-expressed (Zhang et al., 2021). A noise regularization approach was introduced in Zhang et al. (2021) to eliminate such correlation artifacts. However, none of the existing methods have been evaluated within the scope of population-scale scRNA-seq datasets with varying levels of technical artifacts across individual datasets for their reproducibility and stability. Furthermore, their performances in terms of estimating network centrality measures have not been assessed.

Here, we aim to fill this gap by developing Dozer to estimate gene-gene correlations from scRNA-seq data. Unlike existing approaches, Dozer accounts for the noise in the normalized expression and automatically generates a “noise-to-signal” metric as an index to select genes for reliable co-expression analysis. Through large-scale simulations and data-driven computational experiments, we show that Dozer yields robust estimates of gene-gene correlations that are less sensitive to the overall expression levels of the genes and the sequencing depths of the datasets compared to alternatives. Furthermore, Dozer outperforms other methods in estimating network centrality measures such as degree, pagerank, betweenness, eigenvector centrality, and modules. We demonstrate the utility of Dozer in two population-scale scRNA-seq datasets where Dozer constructed co-expression networks have higher proportion of edges validated by external datasets compared to others. These applications further showcase how network centrality measures from personalized co-expression networks can be leveraged to exploit phenotypic variation.

## Results

### Correction factors for correlations estimated from normalized expression data

We consider a biologically motivated hierarchical model to disentangle the biological signal and the measurement error resulting from the sequencing procedure. Let ***g***_*j*_ represent the expression level of gene *j*, ***ℓ*** represent cell sequencing depth, ***X*** represent cell level covariates, e.g., batch labels, mitochondrial percentage denoting the mitochondrial transcript counts as a percentage of the total transcript counts. The observed UMI count of gene *j*, ***Y*** _*j*_, is modeled as

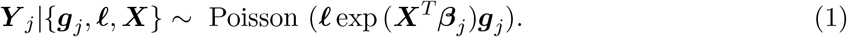

This Poisson measurement model succeeds in capturing variation driven by sampling noise and stochastic technical noise (Sarkar and Stephens, 2021; Choudhary and Satija, 2022) and is particularly well suited for datasets with shallow depths common to population-scale scRNA-seq studies. While an explicit expression model for ***g***_*j*_ is not required for correlation estimation, in our simulation studies, we consider a Gamma prior ***g***_*j*_ ∽ Γ(*v*_*j*_, *u*_*j*_), where *v*_*j*_ and *u*_*j*_ are shape and scale parameters. With this expression model, UMI counts follow the widely used negative binomial distribution (Hafemeister and Satija, 2019; Huang et al., 2018). Let 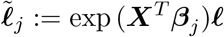 denote the normalizing size factor and 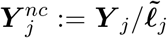 denote the normalized counts. Elementary calculations result in

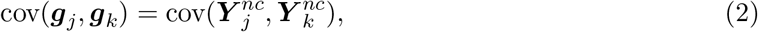

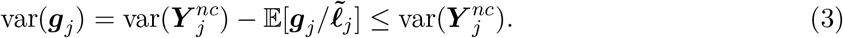

Hence, the magnitude of the correlation of the normalized counts 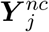 and 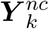 is an underestimate of the true correlation, cor(***g***_*j*_, ***g***_*k*_), of genes *j* and *k*. We use the ratio 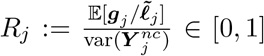 to define a “noise ratio” indicating the *quality* of normalized expression of gene *j*. Quantification of this ratio across a wide range of genes in both simulated and actual datasets indicate that high noise ratio corresponds to low expression and high sparsity (Supplementary Figure S1, Supplementary Section 1.1). This aligns well with the intuition that the sparser the gene expression, the harder to recover the underlying signal. The corrected correlation between the expression values of gene *j* and *k* is then given by

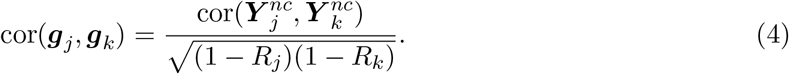

For a given dataset, ***Y*** _*j*_, *j* = 1 …, *G* are the observed UMI counts of the genes. The cell sequencing depth for cell ***ℓ*** is typically approximated with the total number of UMI counts per cell (e.g., in Hafemeister and Satija (2019)), which is further justified with an approximation error 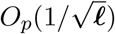 under the Poisson measurement model and probability simplex constraint 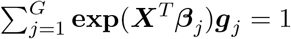 (Zhang et al., 2020a). We used trimmed total UMI counts (Methods), a modification of total UMI counts, to reduce the influence of high expression expression on the estimation, as a default estimator for the sequencing depth. Parameters for the covariates {***β***_*j*_ }_*j*_ are estimated through a Poisson regression. The numerator and denominator of the noise ratio 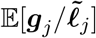, 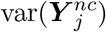, and the gene pair correlation 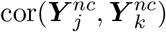 are estimated through the sample mean, variance, and correlation. We denote *S*_*j*_ = 1*/*(1 − *R*_*j*_) as the correlation correction factor for gene *j* and represent cor(***g***_*j*_, ***g***_*k*_) in Eqn. (4) as 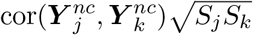. Since the variance of a plug-in estimator 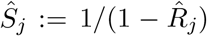 is inflated for genes with high noise ratio (*R*_*j*_ close to 1), we adopt two strategies to stabilize the estimation of the corrected gene-gene correlations: a weighting scheme that allocates higher weights to cells with higher depths in the estimation of gene noise ratio, and a variance reduction transformation to stabilize the gene correction factor estimates (Methods, Supplementary Figure S2).

### Dozer reduces false positives in co-expression network edge estimation

We first designed a data-driven permutation experiment with the Jerber_2021 data (Jerber et al., 2021), leveraging Proliferating Floor Plate Progenitors (P_FPP) cells from 20 donors, each with at least 500 cells, to study the overall false discovery rates of co-expression network construction methods in terms of edge estimation. We also leveraged these computational experiments to investigate the factors that affected the numbers of false positive edges associated with each gene within each method. Given an expression matrix, we randomly split genes into two disjoint sets and permuted the cell ordering in one set so that gene pairs with one gene in each set have zero correlation by construction. We then constructed a co-expression network for gene pairs with one gene from each set using the permuted dataset by each method. Since the networks with permuted data are null networks, all the discovered edges are deemed as false positives.

Quantification of the empirical false discovery rates of each method across the data splitting experiments revealed that Dozer has a smaller false discovery rate than the other methods irrespective of the correlation threshold (Figure 1A, one-sided Wilcoxon rank sum test p-values comparing Dozer vs. the next best method is ≤ 0.077 across all percentile thresholds). We next asked how the numbers of false positive edges of genes varied as a function of the overall expression of the genes and their sparsity i.e., proportion of cells with zero expression for a given gene (high proportion indicating high sparsity) (Figures 1B). Networks constructed by Saver and SCT.Pearson have more false positives among genes with high expression or low sparsity. SCT.Spearman tends to overestimate the number of edges for genes with high sparsity. This appears to be an artifact due to over-smoothing of the normalization procedure and has been observed by others as well (Zhang et al., 2021) (Supplementary Figure S3). In contrast, Dozer, MetaCell, Noise.Reg and bigSCale2 do not exhibit any discernible association between the numbers of false positive edges and the overall sparsity levels of the genes or the overall expression levels of the genes. Next, to elucidate the aggregated effect of the overall mean expression and sparsity levels of the genes, we compared the estimated degrees of the genes, i.e., the total number of edges of a gene, obtained from the networks with the permuted and the unpermuted data. The gene degrees estimated from the networks with unpermuted and permuted data are correlated for methods Saver, SCT.Pearson, and SCT.Spearman (Figure 1C). This indicates that some of the high centrality genes in these networks are driven by the edge identification bias towards genes of certain expression patterns, e.g., high expression genes for Saver, SCT.Pearson, and sparse genes for SCT.Spearman. Dozer, Meta-Cell, Noise.Reg, and bigSCale2 exhibit a uniform behaviour across the gene degrees, with markedly smaller number of false positive edges (an average of 3.4 vs. 9.2, 11.3, and 9.8 in Figure 1) for Dozer compared to MetaCell, Noise.Reg, and bigSCale2, respectively.

**Figure 1:**
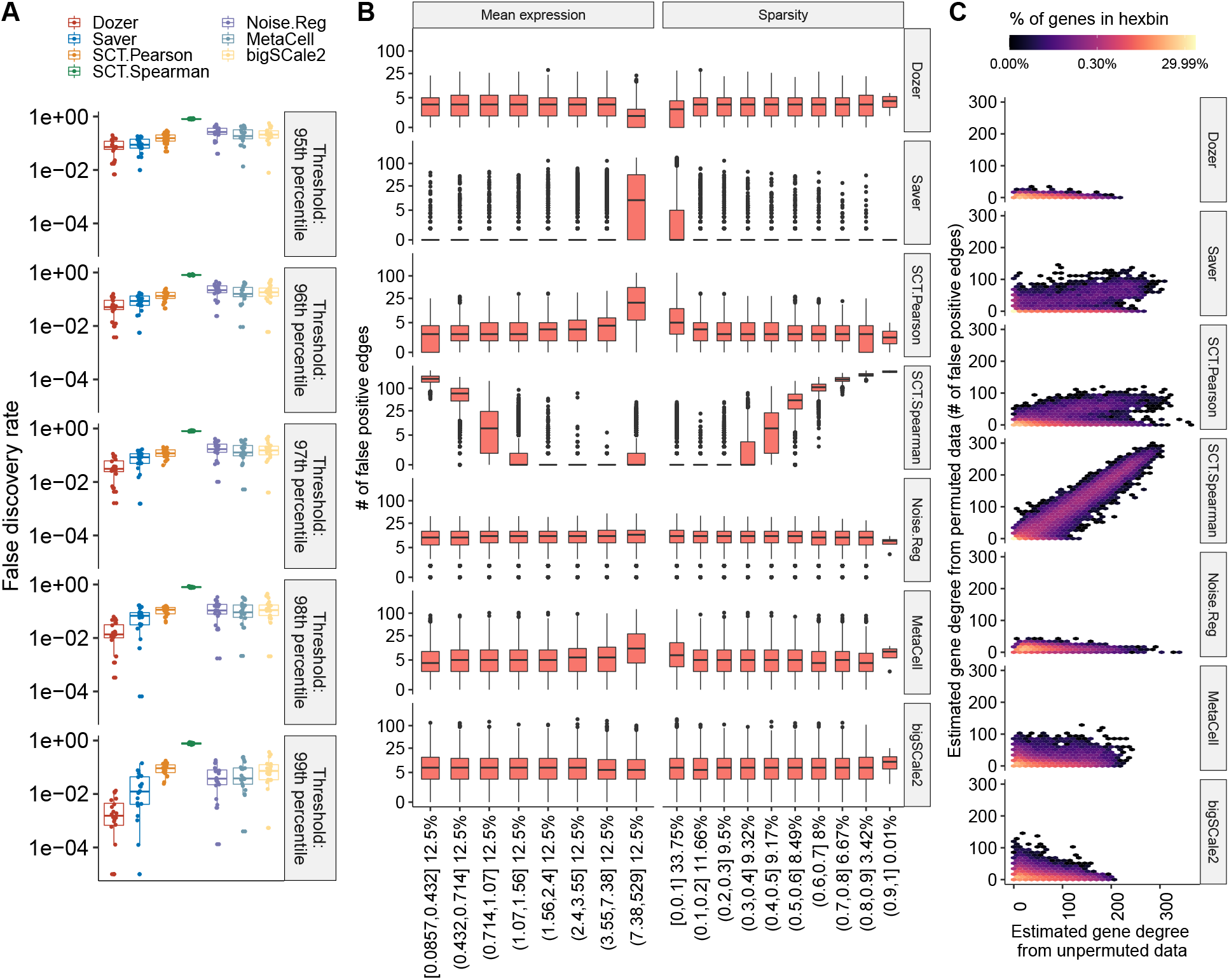
Impact of the overall expression and sparsity levels of genes on false discovery rates of co-expression network edge detection. **(A)** Empirical false discovery rates of the methods across multiple permutation experiments with the Jerber_2021 samples at multiple distinct thresholds set by the percentiles of the absolute values of the estimated correlations. The p-values from one-sided Wilcoxon rank sum test between Dozer and the second best method, Saver, for the five quantile thresholds are 0.077, 0.0014, 4.1 × 10^−5^, 9.5 × 10^−6^, and 0.00016, respectively. **(B)** Numbers of false positive edges of genes stratified by mean expression levels and the proportion of zeros in gene counts (sparsity). Percentages on the x-axis denote the percentage of genes in the expression and sparsity intervals. **(C)** Estimated gene degrees (i.e., numbers of edges connected to a gene) from the co-expression network with the permuted (y-axis) vs. unpermuted (x-axis) data. Co-expression networks are constructed between the two groups of genes resulting from splitting of the genes. Gene degrees are estimated from co-expression networks with the original data (unpermuted, x-axis) and data where cells are permuted for one set of the genes (permuted, y-axis). Since the correlation of genes in the permuted data is zero, the corresponding gene degrees are contributed by falsely detected edges, highlighting the aggregated impact of mean expression levels and proportion of zeros in deriving genes’ overall associations. Results are pooled from multiple permutation replicates across the genes.

### Dozer yields more accurate estimates of gene centrality scores and modules in co-expression networks

Co-expression network construction methods are traditionally benchmarked for their accuracy in detecting individual edges utilizing the Area Under Precision Recall curve (AUPR) and F1 score metrics (Johnson and Krishnan, 2022; Stone et al., 2021; Pratapa et al., 2020). Although inferential analysis of co-expression networks commonly utilize gene centrality measures and modules to prioritize genes and elucidate biological processes (Iacono et al., 2019; Wang et al., 2021), existing methods for estimating co-expression networks from scRNA-seq data are not benchmarked for their performance in estimating these inferential parameters.

To create simulation scenarios that accurately reflect real-world conditions, we relied on Cuomo 2020 (Cuomo et al., 2020), which provides a high sequencing depth per cell (averaging about 530K total counts per cell), to simulate from realistic gene-gene correlation structures. We combined these correlations with marginal gene distributions and sequencing depth estimates obtained from Jerber_2021 (Jerber et al., 2021) to simulate population-scale scRNA-seq data (Methods). In addition to the standard evaluation of network edge recovery, we evaluated each method’s performance in identifying top centrality genes, as defined by a variety of network centrality measures (setting A). We also assessed their abilities to avoid confounding between differential gene expression and differential gene centrality (setting B) and the performance in gene module identification in terms of identifying gene modules in the ground truth network (setting C). The former is important for co-expression analysis of population-scale scRNA-seq data because genes that are differentially expressed between populations might also spuriously exhibit differential network centralities due to the computational biases as observed in the false discovery analysis (Figure 1).

Performances of the methods in terms of edge and high centrality gene identification as a function of noise ratios of the genes (i.e., with four different noise ratio thresholds as {0.6, 0.7, 0.8, 0.9}), the numbers of cells (four different cell sample sizes as {125, 250, 500, 1000}), and average sequencing depths (four different sequencing depths as {1500, 3000, 6000, 12000}) in the first simulation setting (Setting A) are summarized in Figures 2A, B and Supplementary Figures S4, S5. Specifically, Saver tends to outperform the rest in terms of edge identification, while Dozer exhibits superior performance in terms of identifying high centrality genes. The discrepancy in edge and high centrality gene identification performances of Saver can be attributed to its upward bias towards high expression genes, which was also prevalent in the FDR analysis (Figure 1B). Correlation estimation from Saver tends to be larger in magnitude for high expression genes. This upward bias promotes high ranking correlations between high expression genes, leading to more selected edges among these genes (Supplementary Figure S6). Exploring the impact of the noise ratio and the numbers of cells for co-expression analysis yielded that, for all methods, noise ratio and sample size have a large impact on edge identification but much smaller impact on the centrality measures of the genes (Figures 2A, B and Supplementary Figures S4, S5). For the three top performing methods Dozer, Saver, and SCT.Pearson, the AUPR of edges reduces by 47% (F1 score reduces by 38%) when we relax the noise ratio threshold from 0.6 to 0.9 or decrease the numbers of cells from 1000 to 250, while the AUPR of gene centrality is attenuated by only 18% (F1 score reduces by 15%). Visualizing the general trends in left most panels of Figures 2A, B (also Supplementary Figures S4, S5), we observe a clear drop in performance when going from 250 to 125 cells (AUPRs of edge and degree centrality identification are (0.22, 0.58) with 250 cells and (0.12, 0.42) with 125 cells), especially for datasets with low sequencing depths. This indicates the importance of extra caution and additional robustness checks when constructing networks with fewer than 100 cells. Further extension of these simulations by varying the sequencing depths of the cells reveals that the sequencing depth is not a key factor for network construction performance as long as genes with high noise ratio are filtered. However, sequencing depth plays a key role on the numbers of genes retained, hence the size of the network which can be accurately recovered (Supplementary Figure S7).

**Figure 2:**
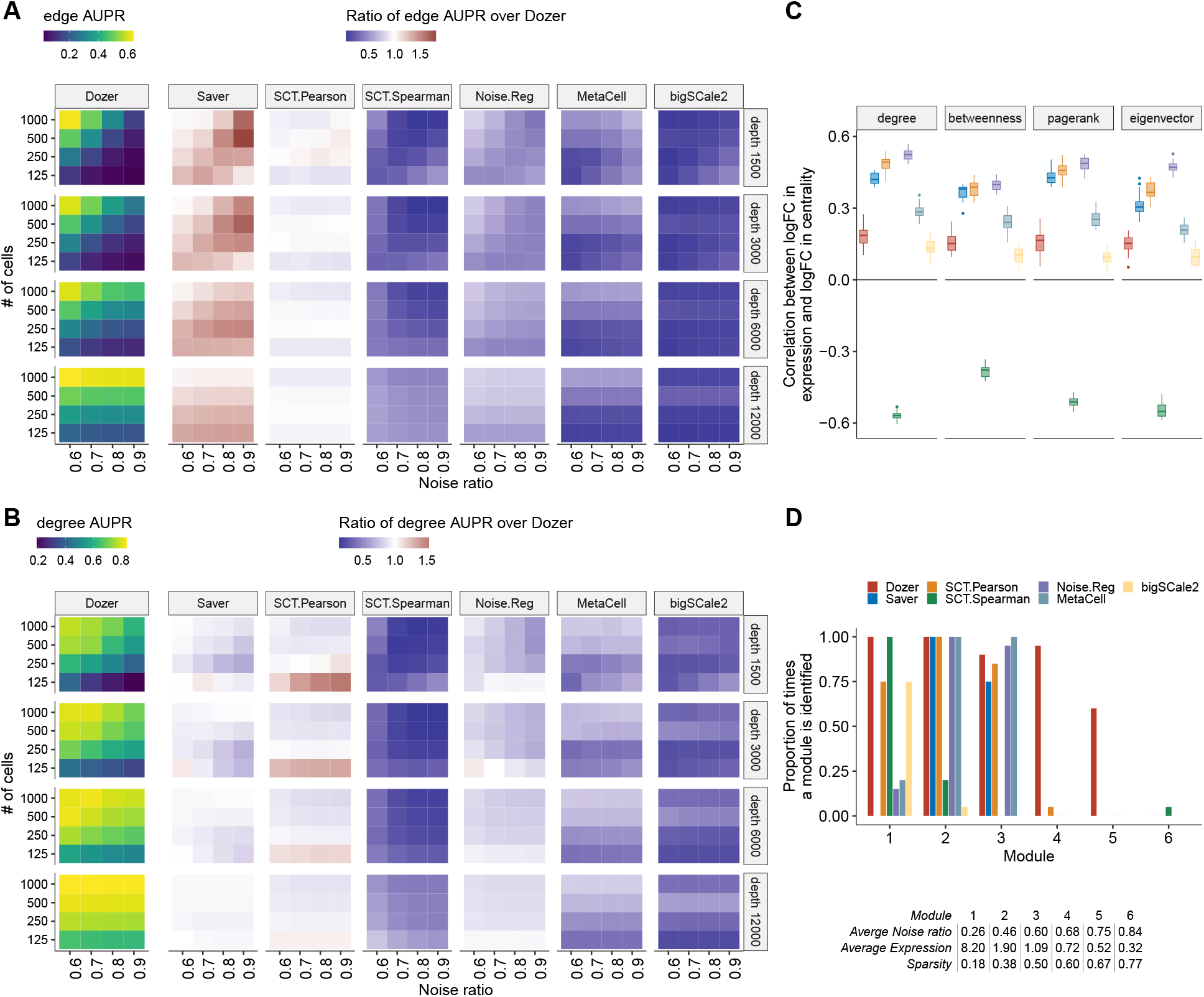
Evaluation for gene centrality and module estimation. **(A-B)** Summary of simulation results in terms of AUPR scores for edge and high degree centrality gene identification. The left most panel depicts the average AUPR scores of Dozer for (A) edge and (B) degree centrality identification as a function of gene noise ratios (x-axis), number of cells (y-axis), and average sequencing depths (rows). The remaining panels highlight the performances of other methods as quantified by the ratio of their AUPR scores over the AUPR score of Dozer. Results for the identification of high centrality genes with respect to pagerank, betweenness and eigenvector centrality are in Supplementary Figure S4. **(C)** Robustness of estimated gene centrality measures against differential expression. Box plots of Spearman correlations between log fold changes (logFC) of expression and centrality across genes are displayed for individual methods. While the two batches of simulated datasets have induced differential expression, they share the same co-expression network structure, hence leading to differential expression but similar centrality measures of the genes across the two batches. **(D)** Proportion of times each module is identified by each method. A gene module was deemed as identified by a method if it has a Jaccard Index overlap of at least 0.5 with WGCNA estimated modules of the method. The table below the barplot provides the average sparsity, expression, and noise ratio of genes in each module.

While Dozer does not assume a Gamma expression model, we further evaluated its robustness under additional data generative settings. Specifically, we compared Dozer with the other state-of-the art methods using `scDesign2` (Sun et al., 2021) (Supplementary Section 1.2). ScDesign2 chooses the marginal distributions of the genes adaptively among a larger set of count distributions (Poisson-Gamma, Negative Binomial-Gamma, Zero Inflated Poisson-Gamma, and Zero Inflated Negative Binomial) and employs a Copula model to generate gene-gene correlations. In experiments with `scDesign2`, Dozer demonstrated robustness to the violation of the Poisson-Gamma assumption and outperformed other methods (Supplementary Figures S9 and S10).

Next, we designed a second batch of simulations (Simulation setting B) to evaluate potential confounding between changes in expression and changes in network centrality of the genes. This type of confounding can lead to inaccurate inference. For example, an increase in expression of a group of genes due to perturbation might be inferred as having interactions with a larger groups of genes due to biased correlation estimation. Each simulation instance contains two datasets generated from the same underlying network (i.e., same gene-gene correlation matrix), but with different gene expression levels controlled by the shape and scale parameters of the Gamma-Poisson distribution. After constructing co-expression networks with each method, we quantified, for each gene, the log fold change (logFC) of centrality between the two datasets in the same simulation instance. We also quantified the logFC of gene expression across the same two datasets for each gene. Figure 2C displays the associations of these two fold changes as quantified by the Spearman correlation across the genes. Since the centralities are estimated from data originating from the same underlying network, methods robust to differences in expression levels are expected not to yield significant associations between logFC of gene centrality measures and expression. We observe that bigSCale2 and Dozer exhibits relatively small correlations between differential expression and centrality measures. In contrast, the logFCs of gene expression and centrality have strong positive associations for Saver, SCT.Pearson, and Noise.Reg, and strong negative association for SCT.Spearman. This result aligns with the upward bias of edges towards high expression genes for methods Saver and SCT.Pearson and the oversmoothing issue with SCT.Spearman observed in the FDR analysis.

Finally, we evaluated the methods in terms of their co-expression module identification performances (Simulation setting C). The ground-truth modules, which were balanced with 80 to 100 genes per module, were set as WGCNA (Langfelder and Horvath, 2008) computational modules with the true network as input to the WGCNA. The table in Figure 2D summarizes the general characteristics of the modules in terms of the sparsity, expression, and noise ratio, calculated by averaging over genes in each module. Each method’s co-expression modules were derived by WGCNA. In a simulation instance, a true module was considered as identified by a given method if one of its co-expression modules overlapped with the true model with a Jaccard Index of at least 0.5 (overall results appeared robust to this choice of cutoff). Figure 2D displays the proportion of times the modules are identified by each method and indicates that modules with high noise ratios (modules 4, 5, and 6 with noise ratios greater than 0.65) are harder to identify for all the methods. Re-markably, Dozer exhibits a robust performance for all the modules with an average noise ratio less than 0.75. None of the methods were able to consistently identify module 6 because the correlation among genes with such high noise ratios are shrank toward 0, leaving these genes as singletons in the network. Saver, Noise.Reg, and MetaCell are also challenged in identifying module 1 despite the lowest noise ratio of this module. A closer investigation revealed that, without correction, these genes tend to have high spurious correlations with the other genes. As a result, genes in module 1 were merged with genes in other modules, which hindered the identification of module 1. Overall, Dozer is more robust in identifying modules with an average accuracy of 0.74 across the modules, 68% higher than the second best method SCT.Pearson.

### Dozer is robust against sequencing depth differences of the scRNA-seq datasets

In large-scale scRNA-seq studies, data are typically generated in separate batches due to logistical constraints, e.g., at different times, laboratories, with different library preparation technologies, and so on. Although this can cause systematic differences in expression between batches and requires correction (Tran et al., 2020) before the data can be analyzed jointly, it further provides an opportunity to evaluate the impact of sequencing depth on estimated network features as the underlying co-expression networks of these batches are realizations from the true co-expression network. While we evaluated the robustness of estimated network features against differential expression with simulation setting B in the previous section, we reasoned that the multiple batches setting can further corroborate our findings with actual data. Figure 3A displays sequencing depths of two biological replicate datasets sequenced in different batches (labelled as pool2 and pool3) from the Jerber_2021 (Jerber et al., 2021) study. Median total read counts in pool2 is 70% of the median total counts in pool3. There is also a clear separation of cells in the two batches in the UMAP (Becht et al., 2019) visualization (Figure 3A). We utilized gene-gene correlations estimated from pool3 (with the higher depth) as the reference to quantify the effect of lower sequencing depth on correlation estimation. Specifically, we evaluated the root mean square error of the absolute correlations (Figure 3B, Method) to quantify the similarity of the correlations from the two batches. Dozer, Saver, and SCT.Pearson yielded lower root mean squares than other methods regardless of the noise ratios of the genes, indicating robustness of estimated correlations against sequencing depth differences of the scRNA-seq datasets.

**Figure 3:**
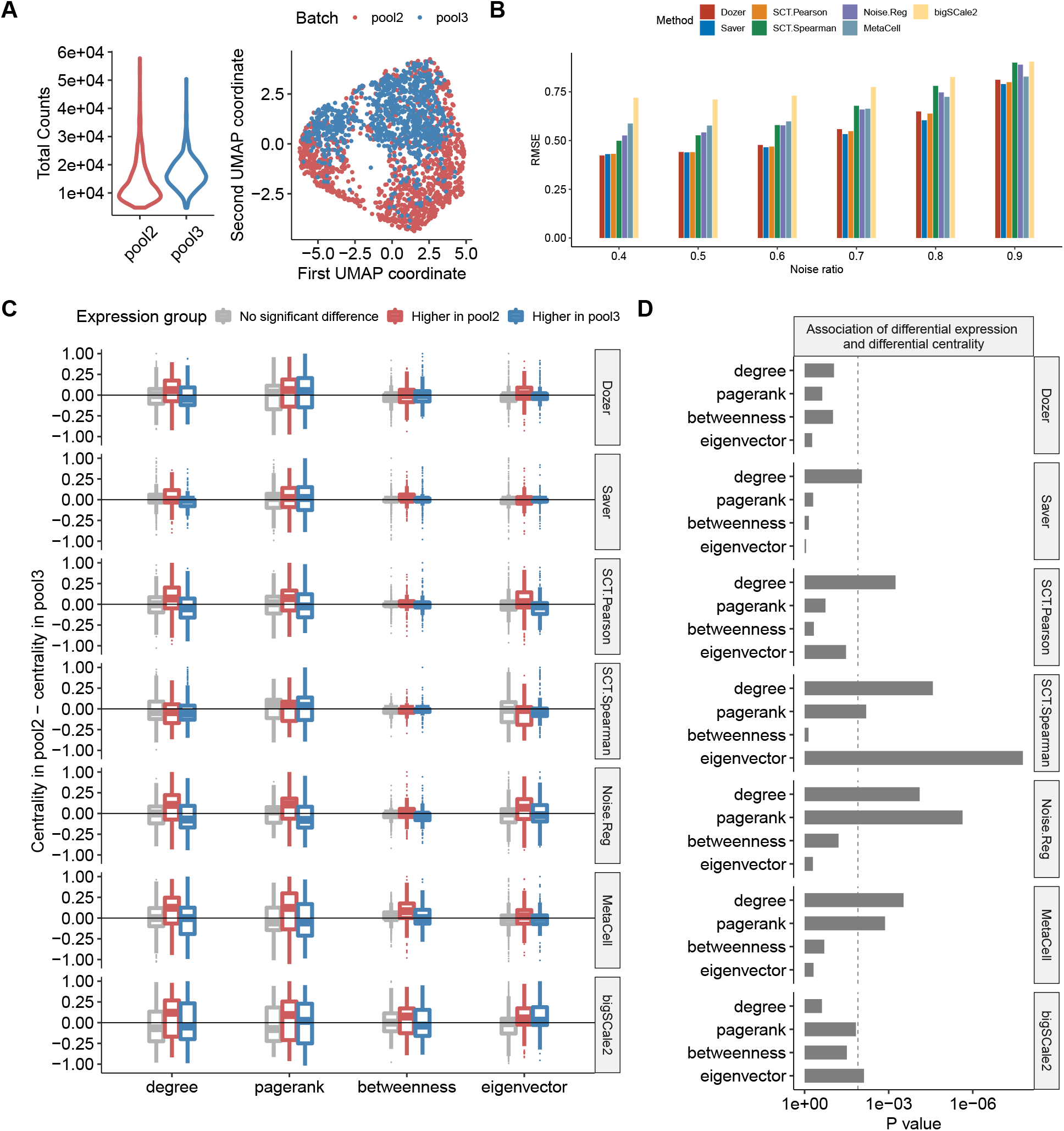
Robustness in co-expression networks against sequencing depth differences of the scRNA-seq datasets. **(A)** Distribution of sequencing depths across cells and UMAP visualization of the cells in biological replicates pool2 and pool3. **(B)** The mean squared error of correlation estimates in pool2 using the higher depth pool3 as the gold standard. **(C)** Genes are separated into three groups as “higher expression in pool2”, “higher expression in pool3”, and “no significant differential expression” using an adjusted p-value threshold of 0.05. For each gene group, the boxplot displays the differences in gene centrality scores between the pool2 and pool3 datasets. Methods robust to sequencing depth differences have centrality differences centered at zero regardless of the gene group. **(D)** P-values from testing the association of differential expression and differential centrality. Dashed line is the Bonferroni corrected p-value threshold of 0.05*/*4.

Next, we considered the impact of expression differences due to batch effects on gene centrality estimates from the co-expression networks. With population scale data, a key downstream analysis is to detect network-level differences associated with phenotypic or genotypic variation (Forbes, 2022). If a network construction method is biased by the actual expression levels of the genes, differential expression will impact the detection of network-level changes. We reasoned that while the differences in sequencing depths or other experimental artifacts would lead to differentially expressed genes between the two batches, robust estimation of gene-correlations should not result in genes with differential network centrality measures. A differential expression analysis with DESeq2 (Love et al., 2014), identified 231(270) genes with significantly higher expression in pool2 (pool3) compared to pool3 (pool2). The choice of DESeq2 ensured that all the methods had access to the same set of differentially expressed genes. We then assessed if a spurious change in expression is carried over to a change in gene centrality measures. This revealed significant associations between differential expression and centrality, including degree, pagerank and eigenvector centralities, for all methods except Dozer (Figures 3C, D). Overall, Dozer provides the most protection against the carry-over effects from changes in expression to centrality.

### Personalized co-expression network analysis of donor iPSC lines under neuronal differentiation identifies genes central to differentiation efficiency

The Jerber_2021 (Jerber et al., 2021) study, with multiple human induced pluripotent stem cell (iPSC) lines differentiating towards a midbrain neural fate, is one of the pioneering population-scale genetic studies with single cell RNA-seq profiling. Neuronal differentiation efficiency score is quantified for each donor iPSC line in the original study as a phenotypic trait. We specifically focused on the Proliferating Floor Plate Progenitor (P FPP) cells to discover genes related to neuronal differentiation efficiency and evaluate the resulting co-expression networks from different methods with external data sources.

For each donor, we constructed a gene co-expression network, by keeping edges with absolute estimated correlations greater than the *x*th-percentile, *x* ∈ {90, 95, 99}, of the all absolute estimated correlations. We first evaluated the edges identified in the co-expression networks with the STRING protein-protein interaction database (Szklarczyk et al., 2021). Across all the methods, a larger fraction of network edges overlapped with the interactions in the STRING database with a higher threshold (Figure 4A), suggesting that the gene pairs with large absolute correlations are better supported by the corresponding protein-protein interactions. Overall, edges from Dozer network had the highest validation rate with the STRING database for all thresholds, with an average increase of 13%, 25% and 56% in the percent validations, for the percentile thresholds of 90%, 95% and 99%, compared to the next best method (one-sided Wilcoxon rank sum test p-values for Dozer vs. the next best method in Figure 4A are 1.1 × 10^−11^, 3.9 × 10^−12^ and 4.1 × 10^−12^, under the three thresholds, respectively).

**Figure 4:**
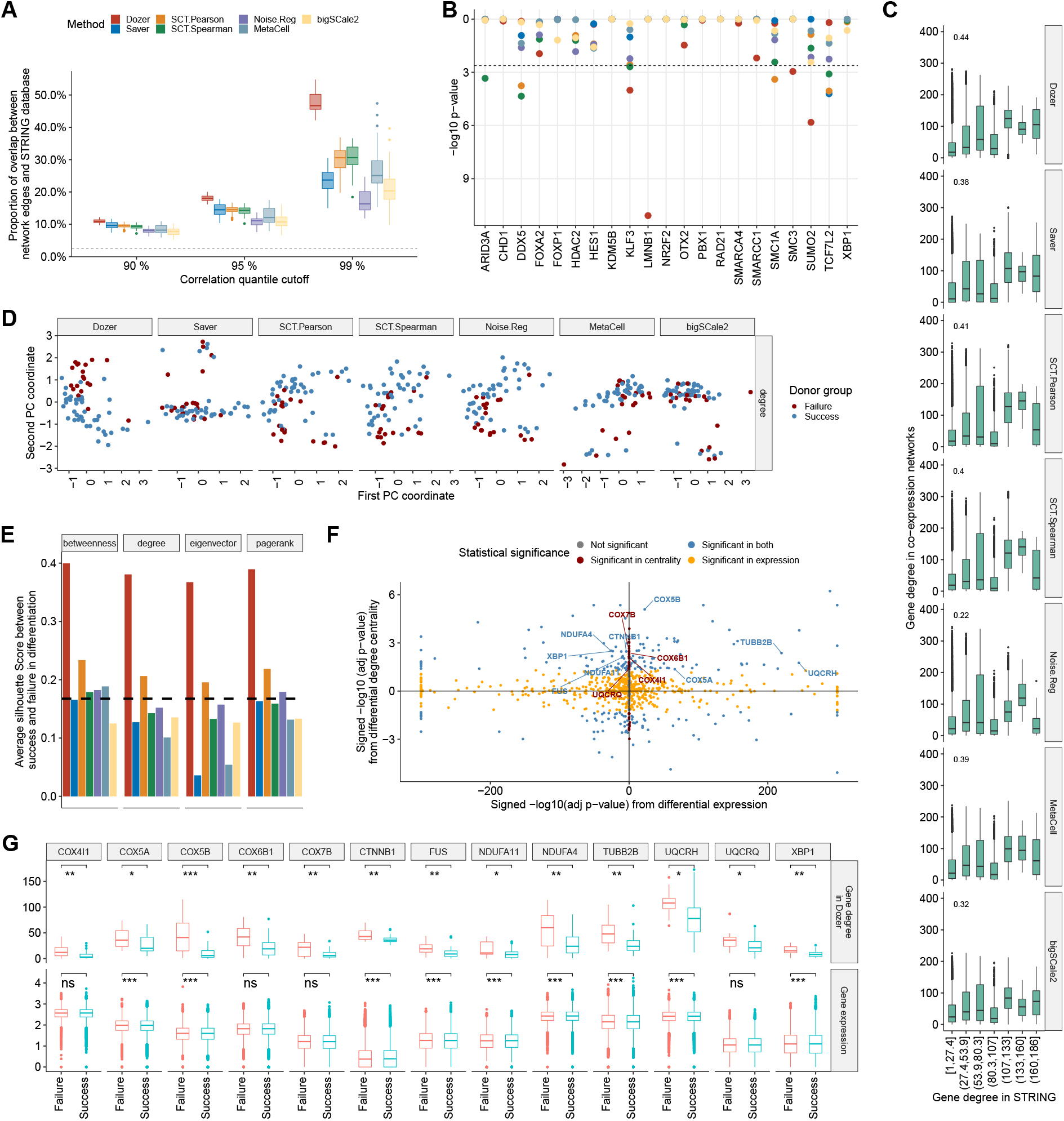
Co-expression network analysis of the Jerber_2021 multiple donor scRNA-seq data. **(A)** The percentage of co-expression network edges validated by the STRING protein interaction database across donors. X-axis denotes the percentile cutoff for thresholding the estimated correlations. Dashed line is the percentage of randomly selected gene pairs validated in the STRING database. The p-values from one-sided Wilcoxon rank sum test between Dozer and the second best method for the three percentile thresholds are 1.1× 10^−11^, 3.9 × 10^−12^, and 4.1× 10^−12^, respectively. **(B)**Transcription factors (TF) enriched in gene co-expression networks of the donors, evaluated using TF-target pairs documented in the hTFtarget database. **(C)** Comparison of estimated gene degrees from co-expression networks with gene degrees in the STRING database. **(D)**Visualization of donor specific networks using the first two principal components of the network degree centralities. **(E)** Average silhouette scores from the first two principal components of the two groups of donors based on their gene centralities in neuronal differentiation. The dashed line represents the average silhouette score based on principal components of donors’ bulkified expression. **(F)** Comparison of differential degree centralities of the genes from the Dozer co-expression networks with their differential expression. X-axis displays the signed -log10(adjusted p-values) from differential expression with positive (negative) values denoting higher (lower) expression in the Failure group. Y-axis denotes the signed -log10(adjusted p-value) from differential degree centrality with positive (negative) values exhibiting higher (lower) centrality in the Failure group. **(G)** Comparison of the degree centralities from the Dozer co-expression networks and expressions of select genes associated with “neurodegeneration” across the donors in the two neuronal differentiation efficiency groups. Significance levels are coded as: ∗: adjusted P-value *<* 0.05, ∗∗: adjusted P-value *<* 0.01, ∗ ∗ ∗: adjusted P-value *<* 0.001.

Next, we utilized hTFtarget (Zhang et al., 2020b), a database of human transcription factor targets, for further validation of the edges connected to transcription factors. Specifically, we tested whether the targets of TFs were enriched among the edge genes of TFs in the co-expression networks (Methods). Four TFs exhibited significant enrichment in the Dozer networks (Figure 4B) while most networks did not yield any enrichment (MetaCell, Noise.Reg, bigSCale2) and Saver, SCT.Spearman, and SCT.Pearson resulted in 1, 4 and 3 TFs enriched for edges by their hTFtarget targets, respectively. Of these, Lamin B1 (*LMNB1*) which is identified only by Dozer, modulates differentiation into neurons (Mahajani et al., 2017). *SMC3* is a member of the cohesion complex, which plays a critical role in regulating changes in chromatin structure and gene expression (Ball Jr et al., 2014). In a study with *Smc3* -knockout mice, reduced cohesion expression in the developing brain resulted in alterations in gene expression, which subsequently caused distinct and abnormal neuronal characteristics (Fujita et al., 2017).

We next turned our attention to gene centrality measures estimated from these co-expression networks. First, to validate gene degrees, we compared the network gene degrees in the co-expression networks and in the networks formed by gene pairs with interactions in the STRING database (Figure 4C). The Spearman correlation between the degrees of the genes in co-expression network and the STRING network is the highest for Dozer, at a level of 0.44. This reinstates that Dozer does not sacrifice accuracy of gene degrees for edge accuracy.

In the Jerber_2021 (Jerber et al., 2021) study, the donors are divided into two groups in terms of their neuronal differentiation efficiency, namely, neuron differentiation failure (neuronal differentiation efficiency *<* 0.2) and neuron differentiation success (neuronal differentiation efficiency ≥ 0.2). We asked whether network gene centrality measures can highlight differences between the two phenotype groups. After estimating a variety of gene centrality measures from each donor’s co-expression network, we visualized the separation of the two donor groups with principal components (PC) and assessed the level of separation with the silhouette score (Rousseeuw, 1987) higher positive values of which indicate good separation between the two groups (Method). Centrality measures estimated from the Dozer co-expression networks exhibit a clear separation between the two neuronal differentiation efficiency groups in the PC plots for all the four centrality measures (Figure 4D, Supplementary Figure S12). The silhouette score associated with Dozer is also the largest, with an average of 0.38 among the four centrality types (Figure 4E).

Centrality measures from co-expression networks can lead to identification of biologically relevant genes that might be missed by standard analysis of differential expression and clustering. To this end, we tested for differences in gene centrality measures of the two phenotype groups and compared the differential centrality and differential expression quantification of the genes (Methods). This analysis identified 51 genes that exhibited differential degree but equal expression between the success and failure groups at an FDR of 0.05 (Figure 4F, Supplementary Figure S13 for the other centrality measures). We performed gene set enrichment analysis separately on KEGG pathways and GO Biological Processes for genes that showed significantly higher centralities in the Failure group (adjusted p *<* 0.05 and logFC(Failure/Success) *>* 0), and the Success group (adjusted p *<* 0.05 and logFC(Failure/Success) *<* 0). We identified a set of 13 genes (COX5A, COX5B, COX4I1, COX6B1, COX7B, CTNNB1, FUS, NDUFA11, NDUFA4, TUBB2B, UQCRH, UQCRQ, XBP1) that exhibited significantly higher centrality in the Failure group and were associated with “Pathways of neurodegeneration”. This set of genes does not appear to be identifiable through differential expression analysis, with four genes exhibiting higher expression in the Failure group, five genes with higher expression in the Success group, and four genes yielding equal expression in both groups (Figure 4F, G). Furthermore, this set of genes overlapped with gene sets enriched in neurodegenerative diseases, such as Parkinson’s disease, amyotrophic lateral sclerosis, Alzheimer’s disease, as well as GO terms related to mitochondrial electron transport (Supplementary Figure S14, S15, S16). The biological relevance of the latter is supported by the growing body of literature that suggests that mitochondria are central regulators in neurogenesis (Brunetti et al., 2021; Arrázola et al., 2019). Mitochondrial dysfunction, especially in the electron transport chain, is responsible for neurodegenerative diseases (Guo et al., 2013; Hroudováet al., 2014; Kausar et al., 2018). We next asked whether bigSCale2, which had been utilized for identifying genes with co-expression network centrality differences (Iacono et al., 2019), could similarly reveal biologically relevant gene groups with differential centrality between the two phenotype groups. While bigSCale2 identified a total of 11 genes with differential centrality across the four centrality measures (with only two genes in the gene set not exhibiting differential expression), this set of genes lacked enrichment for neuronal differentiation GO terms.

### Differential analysis of personalized co-expression networks uniquely identifies a dense innate immune response module in the Alzheimer’s disease diagnosis donors

The Morabito_2021 dataset (Morabito et al., 2021) profiled transcriptome of nuclei isolated from the prefrontal cortex (PFC) of postmortem human tissues from 11 late-stage Alzheimer’s Disease (AD) subjects and 7 age-matched cognitively healthy controls. A co-expression analysis is performed in the original paper with single-nucleus consensus WGCNA (scWGCNA) using both single cell and bulk RNA-Seq data. To directly compare donor-specific co-expression networks with the scWGCNA results, we started with the same set of 1,252 genes that scWGCNA utilized leveraging both the snRNA-seq and bulk RNA-seq data. When using only the snRNA-seq data, 682 of these were filtered either due to their zero expression in one or more donor datasets or high gene noise ratios (Supplementary Figure S17). We primarily focused on oligodendrocytes since this cell type constituted, on average, 60% of donor cells, hence providing a sizable sample per donor.

We first established that donor-specific co-expression networks are biologically sound by a largest-clique, i.e., largest fully connected sub-network, analysis that was validated in the independent snRNA-seq dataset from Nagy et al. (2020) (Supplementary Materials: Section 3, Supplementary Figure S18). Next, to identify sub-networks, i.e., modules, driving the variation of co-expression networks between the AD and control diagnoses, we constructed a “difference network” by first averaging the donor-specific unsigned Dozer networks within the AD and the Control diagnoses separately and then taking the differences of these two averaged networks. Hierarchical clustering of the difference network yielded three gene modules, named Dozer-A1, Dozer-A2, and Dozer-A3 (Figure 5A). Taking advantage of the co-expression networks at the donor level, we asked whether densities (i.e., connectedness) of these modules varied between the two diagnoses and observed that module Dozer-A3, with 57 genes, has significantly higher module density in AD compared to the Control diagnosis (Figure 5B, t-test p-value of 0.0084). We found that the majority of the GO terms enriched in this module are related to innate immune response (Figure 5C). We next explored the expression patterns of the genes in Dozer-A3 (Figure 5D, Supplementary Figures S19, S20, S21). It is evident, especially in the hive plots of Dozer-A3 (Figure 5D, Supplementary Figure S21), that the genes with high degrees are not expressed at high levels. More critically, genes driving the differences in the connectivities of the AD and Control groups are also not differentially expressed. Further investigation into these genes revealed that they tend to be co-expressed in a small group of immune oligodendrocyte cells (immune ODCs; cluster ODC13 in Morabito et al. (2021)) and suggested immune ODCs as the primary source driving the module density differences between the AD and Control groups. This is corroborated by the significant co-expression levels of Dozer-A3 module genes in cells of the donors (Supplementary Figure S22). Furthermore, the association between co-expression of Dozer-A3 genes and whether a cell is an immune ODC cell is stronger among the AD diagnosis compared to Control (Supplementary Figure S23). To further validate that immune ODCs are driving the discovery of Dozer-A3, we repeated the difference network analysis with Dozer with a restricted ODC cell population excluding the immune ODCs and observed that immune related gene module Dozer-A3 was no longer discoverable (Supplementary Figure S24). In addition, none of the detected modules from this restricted population showed differences in connectivity between the Control and AD groups. This further corroborated the driver role of immune ODCs in the original Dozer-A3 module.

**Figure 5:**
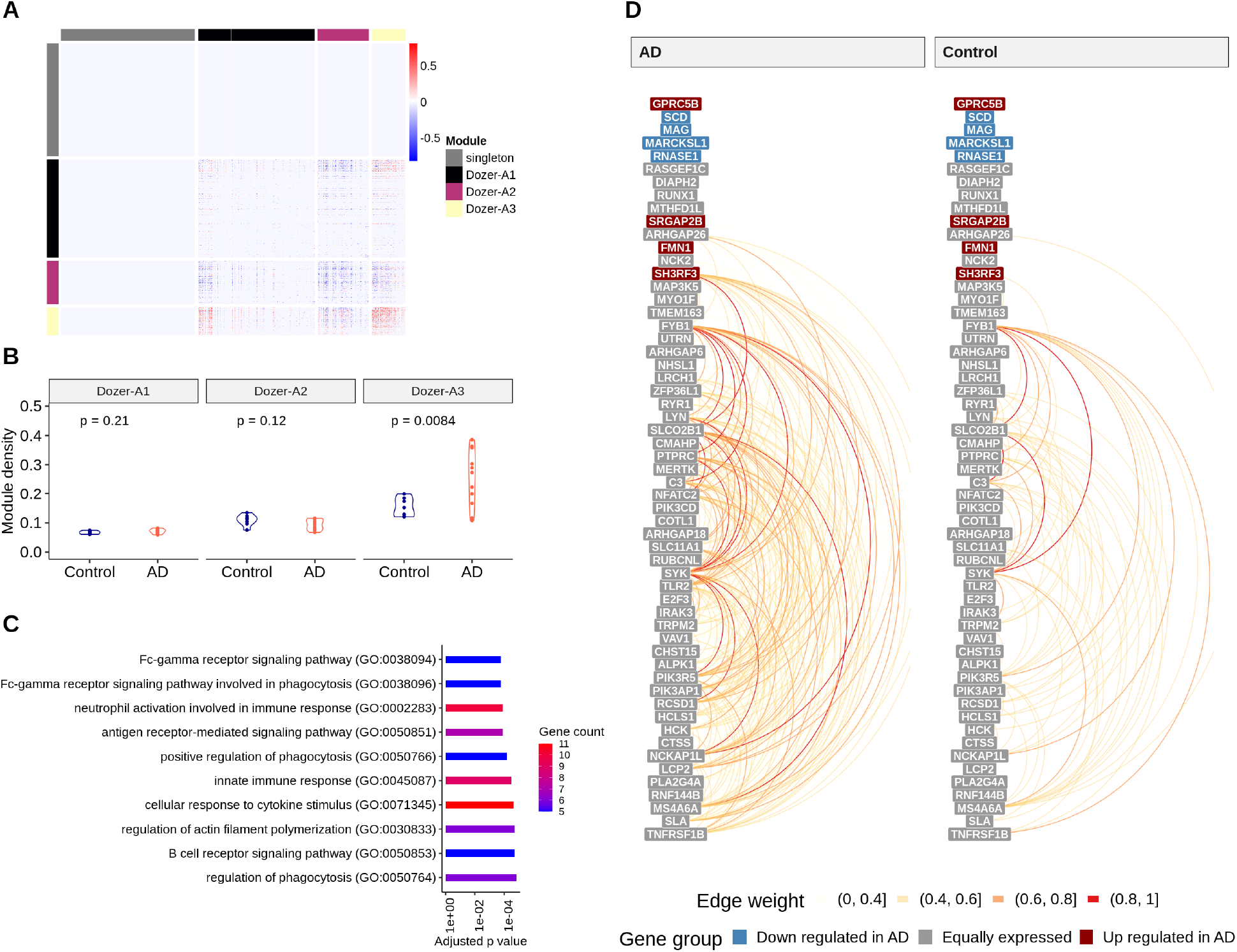
**(A)** Heatmap of the “difference network” (average of AD networks - average of Control networks). Genes are order by gene modules (singleton, Dozer-A1, Dozer-A2, Dozer-A3), which are obtained by hierarchical clustering on the “difference network” (Method). **(B)** Violin plots of module densities for AD and Control donor networks. Module density is a function of the average absolute correlation of gene pairs in the given module. **(C)** Barplot of GO enrichment terms for module A3. **(D)** Hive plot visualization of module Dozer-A3 in AD and in Control groups. Genes are ordered from high (top) to low (bottom) by their average expression across all donors. Genes are further divided into three groups as “Down regulated in AD”, “Up regulated in AD”, and “Equally expressed” according to their differential expression status between the Control and AD diagnoses. The arcs between the genes in this linear layout depict the edges in the average networks of AD (left) and Control (right) donors, with colors representing edge weights.

We next performed differential centrality analysis for degree, pagerank, betweenness, and eigenvector centralities to infer the key genes in the Dozer-A3 module. This revealed that *TLR2*, which is a primary receptor for Alzheimer’s amyloid beta peptide to trigger neuroinflammatory activation (Liu et al., 2012), has significantly higher degree centrality in AD than in Control (Supplementary Figure S25). In the broader context, innate and adaptive immune responses play a key role in the pathological processes of Alzheimer’s disease as well as other neurodegenerative diseases (Shi and Holtzman, 2018). Recent studies have shown that oligodendroglia becomes immune reactive in a mouse model of multiple sclerosis (Kirby et al., 2019) and in human iPSC-derived oligodendrocytes from Parkinson’s disease and multiple system atrophy patients (Azevedo et al., 2022), providing further support for the discovery of this module by Dozer.

Repeating the same type of difference network analysis with the other methods revealed that Dozer, Saver, and SCT.Pearson are the only three network construction methods that unearthed co-expression modules with densities significantly correlated with diagnoses and significantly enriched GO terms (Supplementary Figures S26 - S31). The three modules that exhibit changes in densities between diagnoses are Dozer-A3, Saver-A3, and SCT.Pearson-A3. The three modules from independent methods are supportive of each other, as they share 23 genes (Supplementary Figure S32) and the common set of genes are significantly enriched for GO term “innate immune response” (Supplementary Figure S33).

Finally, we asked whether the association of the diagnosis with the co-expression networks of innate immune response genes is also revealed with the scWGCNA analysis (Morabito et al., 2021) that combines data from all donors to construct a single co-expression network. While Morabito et al. observed that the cell-type composition of late stage AD shifted towards more immune oligodendrocyte cells, the eigengene expression of their immune response related gene module (OM3) within oligodendrocytes is not correlated with the AD diagnosis in the scWGCNA analysis (Figure 8 of Morabito et al. (2021)). scWGCNA, as a consensus clustering approach, utilized external bulk RNA-seq data from UCI (Morabito et al., 2021) and ROSMAP (Mostafavi et al., 2018) cohorts to mitigate the issue of sparsity in expression data. Nevertheless, the inclusion of these external data may have obscured signals that are unique to single-cell RNA-seq. To elucidate the factors that contribute to the differences in findings between scWGCNA and personalized gene co-expression networks, we re-implemented scWGCNA on dataset Morabito 2021, following the gene filtering procedure used in Dozer to exclude genes with high sparsity and noise ratios. Our implementation considered two comparable variants of scWGCNA as: (1) scWGCNA-I: data was combined from donors within a diagnosis group to construct diagnosis-specific co-expression networks, then the consensus modules were computed; (2) scWGCNA-II: a difference network between the diagnoses from diagnosis-specific co-expression networks was constructed and modules on the difference network were inferred. scWGCNA-I follows the prescribed scWGCNA pipeline of (Morabito et al., 2021) with a small modification by using diagnosis-specific networks, while scWGCNA-II is more similar to our difference network analysis for personalized networks and it aims to detect modules showing differences in connectivity between diagnoses. A key difference of both of these implementations from our approach is that, they pool the donor data before constructing networks; hence, the resulting networks are not at the individual donor-level. In these analysis, scWGCNA-I detected three co-expression modules (scWGCNA-brown, scWGCNA-blue and scWGCNA-turquoise) and left a large group of genes as singletons, i.e., deemed as not forming a co-expression module (scWGCNA-grey) (Supplementary Figure S34). Eigengene analysis of these modules revealed only scWGCNA-brown module as weakly associated with the AD diagnosis. Furthermore, innate immune response genes, discovered in our personalized networks analysis, appeared as singletons, prohibiting them to be identified as enriched within a module. This can be explained by scWGCNA-I’s intrinsic focus on modules common to both diagnoses, as a result of which it loses power to detect module-level differences between the diagnoses. scWGCNA-II builds modules on the difference network between AD and Control diagnoses. One of its eight co-expression modules (scWGCNA-A4) harbors the innate immune response genes. However, expression of none of the module eigengenes of scWGCNA-II has significant association with the AD diagnoses (Supplementary Figure S35). In fact, visualization of the Dozer-A3 genes within the scWGCNA networks of AD and Control groups does not exhibit any discernible differences between the two (Supplementary Figure S36). scWGCNA-II pools cells from all donors and loses the ability to directly test differences in module connectivity between diagnoses through module density. This further reinstates that, while module eigengene expression is representative of module gene expression level, when the differences in the expression levels between the diagnoses are small, it might hinder the association of the module with the diagnosis. Constructing personalized co-expression networks as we do with Dozer enables a formal testing framework for downstream association analysis with the modules.

## Discussion

Excess sparsity and measurement error in scRNA-seq datasets distort gene-gene correlation estimation, introducing downward bias for genes with low expression and in low depth datasets. The distortion of estimated correlations has a major influence on the construction of co-expression networks, by uplifting high expression genes to be network hub genes and confounding differential expression with changes in co-expression networks. Dozer, built on a Poisson measurement model, provides correction for gene-gene correlation estimates and offers a gene-specific noise ratio score to reliably filter genes for co-expression network analysis. In our analysis, no restrictions were put on the expression model, i.e., distributional assumption on ***g***_*i*_ in eqn. (1), except for simulation purposes. A large variety of observation models, including Negative Binomial (Huang et al., 2018; Love et al., 2014), Zero-inflated Negative Binomial (Risso et al., 2018), and other flexible models used in (Wang et al., 2018), can be accounted for by combining Poisson measurement error model with Gamma, Point Gamma, or Point exponential family expression models (Sarkar and Stephens, 2021). Although the Poisson measurement model usually suffices in practice, in the cases where a more complex measurement model is more adequate, a gene correction factor should be derived accordingly to adjust for the underestimation of absolute corrections due to measurement error.

For network validation, instead of solely focusing on network edges, i.e., highly correlated gene pairs, we also validated a broader set of network features used in downstream inference, including gene modules and gene centrality measures. We observed a marked discrepancy between edge accuracy and centrality/module accuracy. Most notably, our computational experiments revealed that an upward bias towards high expression genes could lead to an increase in edge accuracy; however, this comes at the cost of decreased accuracy for identifying high centrality genes among low expression genes. This broadly suggests that benchmarking studies of co-expression networks focusing solely on network edges may be inadequate. Both data-driven simulation and computational experiments demonstrated Dozer’s superior performance in mitigating the sequencing depth differences between donor-specific datasets and faithfully preserving the co-expression network structures both at edge, centrality, and module levels, in the presence of expression differences due to technical reasons.

Construction of donor-specific networks enables exploring association of multiple classes of network features with phenotype or genotype information. In contrast, with the existing scWGCNA analysis that pools individual level data before network construction, only the expression of eigengenes of modules can be associated with subject-level information. The Morabito_2021 reanalysis showcased how constructing donor-level co-expression networks can discover a module of genes with significantly different module density between the diagnosis groups even when these genes are co-expressed only in a small sub-population of the cells and are not differentially expressed between the diagnosis groups. Similarly, in the analysis of Jerber_2021, we identified a group of neurodegeneration associated genes with differences in network centrality between the donors that succeeded or failed in neuronal differentiation. However, these genes were not differential expressed between the two phenotype groups further highlighting that donor-specific co-expression networks offer an opportunity to quantify a broader set of network traits.

In conclusion, Dozer is tailored for co-expression analysis with population-scale scRNA-seq datasets and enables further downstream analysis such as network differences between different phenotypic groups with the constructed individual co-expression networks. We envision that similar differential analysis might be of interest between co-expression networks of different cell types. Alternatively, one can formally test whether co-expression networks of different cell types are the same by using recent theoretical developments on testing of large dimensional correlation matrices (Zheng et al., 2019). Additionally, we expect that Dozer-derived co-expression networks when combined with genetic information can facilitate network QTL analysis.

## Methods

### Gene-gene correlations in the Poisson measurement model

Let random variable ***g***_*j*_ represent the expression level of gene *j*, ***ℓ*** represent cell sequencing depth, ***X*** represent cell level covariates. The UMI count of gene *j*, ***Y*** _*j*_, follows a Poisson measurement model (Sarkar and Stephens, 2021),

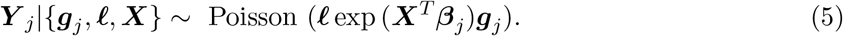

The correlation between the expression levels of gene *j* and *k*, cor(***g***_*j*_, ***g***_*k*_), is the signal we aim to recover for quantifying co-expression between genes *j* and *k*.

Let 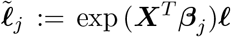 denote the size factor per cell and 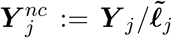 represent the nor-malized counts. Under the conditional independence of the observed UMI counts given the true expression levels and cell sequencing depths

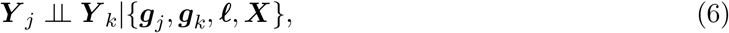

we have

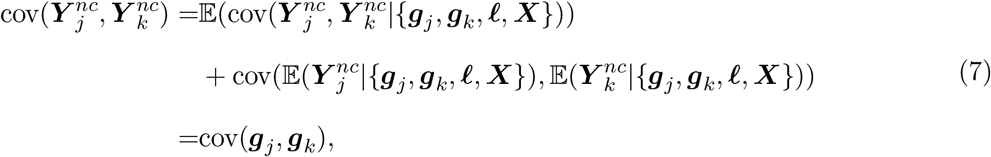

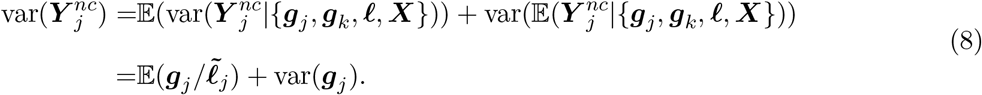

Due to the inflation in the var 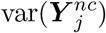 compared to the true variance var(***g***_*j*_), the proxy cor 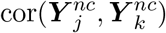 is an underestimate of cor(***g***_*j*_, ***g***_*k*_) in terms of its magnitude. The deviation between the true expression variance of gene *j* and the variance of normalized expression depends on the ratio

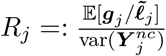 that we denote as gene *j*’s “noise ratio” indicating the quality of gene *j*’s normalized expression. Let 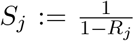, then the expression correlation of genes *j* and *k* can be represented through the following equation with the correction factors:

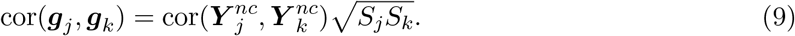

### Estimation of the gene correction factors in the Poisson measurement model

Given the UMI counts ***Y*** _*j*_ and sequencing depths of the cells ***ℓ***, we fit the Poisson measurement model in eqn. (5) and estimate {***β***_*j*_ }_*j*_ via a Poisson generalized linear model

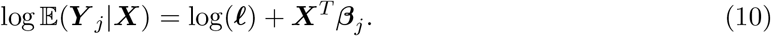

The cell size factor 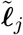 is estimated by plugging in 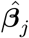 as 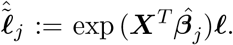. The numerator and denominator of the noise ratio are estimated through sample mean and variance with weights ***w*** as

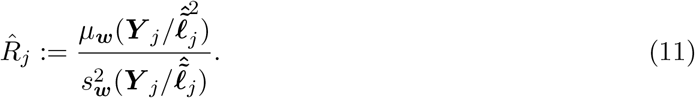

Similarly, the correlation of normalized counts is estimated through sample correlation

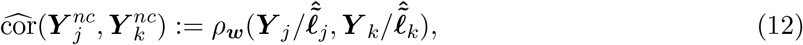

where *μ*_***w***_(·), 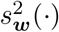, and *ρ*_***w***_(·, ·) denote weighted sample mean, variance, and correlation with weights ***w*** = ***ℓ***, to account for the heteroscedasticity of ***Y*** _*j*_*/****ℓ*** conditional on ***ℓ***. Without loss of generality, the setting with ***β***_*j*_ = 0 provides further insights into this apparent heteroscedasticity. We express the var(***Y*** _*j*_*/****ℓ***|***ℓ***) by utilizing the law of total variance

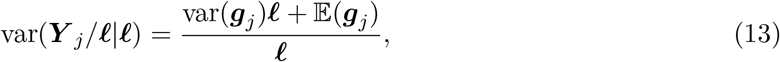

and observe that the conditional variance of ***Y*** _*j*_*/****ℓ*** given ***ℓ*** decreases with depth ***ℓ***. The weight, inspired by the weighted least squares approach, is proportional to ***ℓ****/*(1 + ***ℓ***var(***g***_*j*_)*/*𝔼(***g***_*j*_)). The second term ***ℓ***var(***g***_*j*_)*/*𝔼(***g***_*j*_) in the denominator of the weight is the over-dispersion parameter of gene *j*. To avoid the extra variability introduced by estimation of over-dispersion, we use ***ℓ*** as weights for cells.

The correction factors *S*_*j*_, *j* = 1, …, *G*, cannot be reliably estimated directly by plugging in corresponding 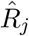 because the resulting *plug-in estimator* can be arbitrarily large for sparse genes with noise ratios close to 1. To robustly estimate the gene correction factor *S*_*j*_, we opted to balance the bias and variance by shrinking the estimator towards zero when its variance is large. Given a plug-in estimator 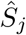 of correction factor *S*_*j*_ with the estimated noise ratio as

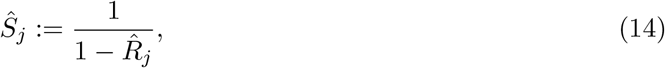

we obtain an initial plug-in estimator of the variance of 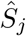 as

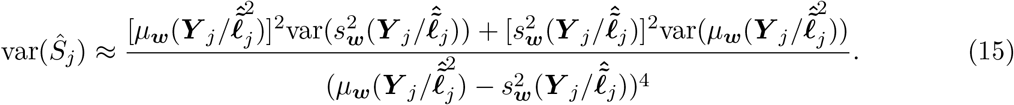

Since the variance of 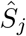 increases with its mean, we consider a penalized correction factor and obtain a *shrinkage estimator* as

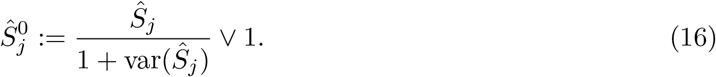

Finally, to borrow information across genes, we fit a local polynomial regression of 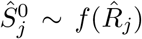 with the R function loess. The fitted function is naturally unimodal, because for genes with noise ratio close to 1, 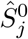 is close to 1. However, the true value of *S*_*j*_ increases with *R*_*j*_. To turn it into a monotone function, we set the turning point *r*_0_ = arg_*r*_ max 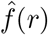, where 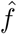 is the estimated local polynomial regression function, and set the final estimate that we refer to as *truncated shrinkage estimator* 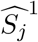as

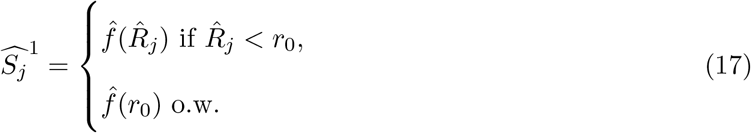

### Sequencing depth estimation

Sequencing depth in cells is typically estimated by the total UMI count per cell (Vallejos et al., 2017). Normalizing gene counts by the total UMI counts imposes a “sum to 1” constraint on the normalized expression. This constraint can result in negative correlations between highly expressed genes. For instance, in a simulation setting without any correlation between gene pairs, this simple normalization process induced a negative correlation of -0.07 between pairs of genes which accounted for 10% of the total UMI counts (Supplementary Figure S8). Prior research has suggested approaches for clipping influential genes, such as the “median of ratios” estimator proposed by Love et al. (2014) and “trimmed mean of M values” of Robinson and Oshlack (2010). Following these ideas, we consider “trimmed total UMI count” as an estimator, designed to mitigate the influence of highly expressed genes. First, we compute the expression proportion of each gene across all cells. These proportions serve as gene weights. Subsequently, we normalize the UMI counts for each gene, ensuring an average value of one. We then set a threshold for gene weights to prevent any single gene from dominating the sequencing depth estimation (e.g., empirically set as 0.02 as this reduced the apparent negative correlation to -0.01 in the simulation setting of Supplementary Figure S8). By limiting the weights of highly expressed genes to the threshold value, we effectively trim their contributions. Finally, we compute the trimmed total UMI count for each cell by summing the weighted UMI counts across all genes as its estimated sequencing depth.

Next, to determine whether a global normalization factor (i.e., same for all the genes) is sufficient for normalization, we perform a diagnostic analysis. Global normalization factors are used to account for a presumed count-sequencing depth relationship that is consistent across all genes. However, when genes of different expression levels grow disproportionately with sequencing depth, normalization through global scale factors can result in over-correction for weakly and moderately expressed genes (Bacher et al., 2017). As part of the diagnostic process, raw gene counts and trimmed total UMI counts are divided by their mean to achieve an average value of one. Then, for each gene *j*, scaled gene counts 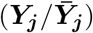 is regressed on scaled trimmed total UMI counts 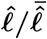 as

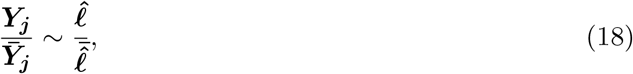

and the slope is estimated from this regression using cell-level data. The diagnostic plots visualize (i) the distribution of these estimated slopes of genes across multiple expression groups, and (ii) distribution of the correlations between the expression of genes normalized with the trimmed total UMI counts. Proportionate growth yields regression slopes around 1 and correlations near 0, while disproportionate growth causes regression slopes and correlations to diverge from these values. When disproportionate growth is detected for a given dataset, Dozer adopts a regression-based gene-specific cell size factor approach for sequencing depth adjustment. This involves first grouping genes into K bins based on their raw mean expression quantiles. Next, the trimmed total UMI counts of each gene group, denoted as 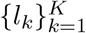, are used as regressors in a Poisson regression to facilitate gene-specific adjustment for sequencing depth. More specifically, for gene *j* with UMI counts ***Y*** _*j*_, we conduct a Poisson regression analogous to eqn.(5) while replacing the global cell size factor *ℓ* with a set of covariates ***L***,

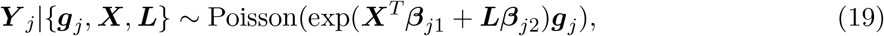

where ***X*** represents cell level covariates as before (e.g., batch labels, percentage of mitochondrial genes), and ***L*** = [log *l*_1_, …, log *l*_*K*_] denotes the design matrix that harbors the trimmed total UMI counts of each gene group for each cell. *K* is chosen incrementally by starting from 1 and gradually increasing it up to 10 until the mode of the correlation between normalized expression and trimmed UMI drops below 0.1 for all gene groups. The estimated cell size factor for gene *j* is then given by

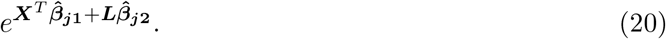

An illustration of how these diagnostics plots and the resulting gene specific size factors work are provided in Supplementary Figure S11 for two donors from the Jerber_2021 dataset. Right panels of the Supplementary Figures S11A and B display the density of correlations between expression normalized with the gene specific cell size factors and trimmed total UMI counts across all the genes. These plots highlight how the apparent correlations between the global size factor and expression normalized with the global size factor are reduced when gene specific size factors are adopted.

### Simulations

Three simulation settings were considered to evaluate the performances of the co-expression network construction methods in recovering network edges and gene centrality measures (setting A), reducing false discoveries in differential network centrality measures in the presence of differential expression (setting B), and module detection (setting C). The following base procedure was used in settings (A-C) to generate the gene expression *Y*_*ij*_ for gene *j*, = 1, …, *G*, in cell *i, i* = 1, …, *N*. First, *G* dimensional relative gene abundance vector ***g*** was generated through a multivariate Gamma distribution, with shape parameters ***v***, scale parameters ***u***, correlation matrix **Σ**. The sequencing depth *ℓ*_*i*_ of cell *i* was simulated from a log normal distribution Lognormal(*l*^*ℓ*^, *s*^*ℓ*^). The UMI count for each gene *j* was sampled independently from Poisson distribution for cell *i* as *Y*_*ij*_ ∽ Poisson (*ℓ*_*i*_*g*_*ij*_). Under this simulation framework, **Σ** specifies the co-expression structure of the genes.

To ensure that these simulations yield data with high fidelity to actual population-scale scRNA-seq data, Cuomo_2020 scRNA-seq data (Cuomo et al., 2020), which offers high sequencing depth per cell (averaging approximately 530K total counts per cell) was utilized to generate realistic gene-gene correlations. These correlations were then combined with marginal expression count distributions from Jerber_2021 (Jerber et al., 2021). Realistic datasets similar to those of Jerber_2021 and Morabito_2021 were simulated by modulating the library sizes. The shape and the scale parameters *v*_*j*_, *u*_*j*_ for gene *j* were estimated by:

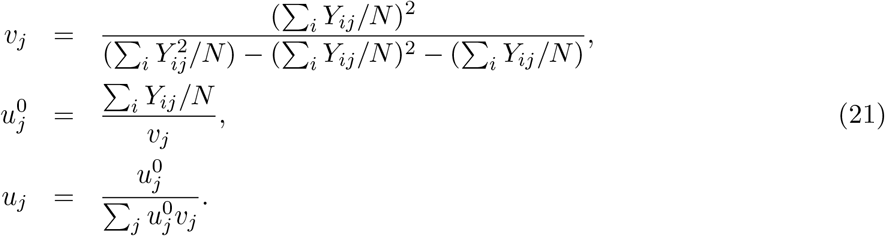

The scaling in eqn. (21) for the scale parameter ensures that the total counts in cell *i* is approximately equal to its sequencing depth *ℓ*_*i*_. The parameters (*l*^*ℓ*^, *s*^*ℓ*^) of the sequencing depth distribution were estimated by fitting a log normal distribution to the total read counts. The correlation matrix **Σ** was estimated through SCTransform normalized expression matrix of the dataset Cuomo 2020.

Further details on the parameters of the simulation settings A, B, and C are as follows.

#### Setting A

This setting considered four average sequencing depth levels (1500, 3000, 6000, 12000) and four sample sizes (125, 250, 500, 1000 cells). For each depth and sample size combination, ten simulation replications were generated from the base model described above. When generating co-expression networks, four noise ratio thresholds, namely, 0.6, 0.7, 0.8, and 0.9, were used for gene filtering. Gene-pairs with top 1% of absolute correlations and genes with top 10% gene centrality measures were set as true edges and true high centrality genes. The same quantiles were applied to estimated networks for AUPR and F1 score calculations.

#### Setting B

For each simulation instance out of 20 replications, two datasets with the same gene-gene correlation structure, but different gene expression levels were generated by reshuffling the Gamma shape and scale vectors. Gene-pairs with top 1% of absolute correlations were used to set the network edges. Both the log2 fold change (logFC) of gene expression and the logFC of centrality were computed across the two datasets of the simulation instance. To avoid taking log of zeros, 1st percentile of positive centrality values across all genes was added to centrality values before taking the log. Spearman correlation between logFC in expression and logFC in centrality was computed to assess the impact of differential expression on detecting spurious differential centrality.

#### Setting C

Across 20 simulation replications, six gene modules, with balanced module sizes of 80 to 100 genes and large differences in average gene noise ratios, were simulated with a block diagonal correlation matrix, with 6 gene modules and 2,434 singletons. Module detection was carried out with WGCNA (with default parameter settings) that took as input the estimated correlation matrices to generate modules. Module detection performances were quantified with the Jaccard index between the inferred and the true gene modules after excluding singletons. If a true gene module had a Jaccard index of 0.5 or larger with an inferred gene module, it was deemed as detected for the purposes of quantifying the empirical probability of detecting gene modules.

### Computation of network metrics and gene centrality measures

#### Thresholding correlations for edges and network centrality measures of genes

Unless otherwise specified, hard-thresholding was adapted for keeping edges between the genes in the co-expression networks and labeling genes based on their network centrality measures. Genes *i* and *j* were set to be connected with an edge in the co-expression network if the absolute value of their estimated expression correlation was larger than threshold *τ*. *τ* was set using the percentiles of the absolute values of the correlations as specified in the analyses throughput the paper.

#### Centrality measures in co-expression networks

Four centrality measures, namely, degree, pagerank, betweenness, and eigenvector centrality, were considered to quantify the “importance” of a gene in the co-expression networks. These were computed using the following functions from the `R` package `igraph: degree, page rank, betweenness`, and `evcent`.

#### Module

Module identification from co-expression networks was performed through the `R` package `WGCNA` (Langfelder and Horvath, 2008) with the default parameter settings with an exception for the difference networks. The canonical WGCNA module identification pipeline starts out with unsigned co-expression similarity measures between nodes/genes as weights. However, the edge weight between two genes in the “difference network”, with positive and negative values, represents whether the association of the genes is higher in AD or in Control rather than a similarity between the two. To accommodate this, we considered a network transformation while still keeping the hierarchical clustering and dynamic tree cut components of the WGCNA pipeline. Specifically, we let 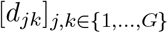 denote the edge weight between genes *j* and *k* in the “difference network”. Then, 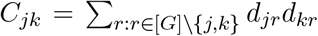 defines a connectivity similarity for genes *j* and *k*. A distance between nodes *j* and *k* can be obtained as *D*_*jk*_ = max 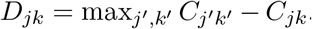. Finally, the gene modules of the “difference networks” are inferred by hierarchical clustering and dynamic tree cut algorithm (`R` function `hclust` and `cutreeDynamic`) using the distance matrix with entries *D*_*jk*_ and the default parameter settings.

#### Module density

Module density, *D*(*M*), measuring the average connectivity of genes within a module was calculated as follows for module M with n genes: 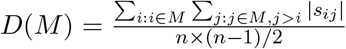, where *s*_*ij*_ ∈ [0, 1] is the absolute correlation between gene *i* and *j*.

#### Largest clique

The largest clique was computed using function `largest cliques` from the `R` package `igraph`.

#### Associating diagnosis with module eigengene expression from the scWGCNA analysis

We used a linear mixed model (22) implemented through `R` package `lmer` to assess the association between module eigengene expression and diagnosis. In equation (22), *y*_*i*_ represents expression of eigengene of metacell *i*. Variables “Diagnosis”, “Sex” and “Age” are fixed effects. “SampleID” was used as a random effect, because metacells in scWGCNA analysis are constructed within samples of individual donors.

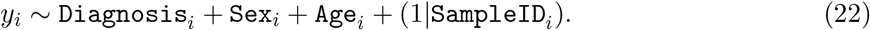

#### Gene set enrichment analysis

Gene set enrichment analysis for gene modules and differential centrality genes is conducted through package `enrichR` (Kuleshov et al., 2016).

#### Silhouette score

Silhouette score (Rousseeuw, 1987) is used to measure the consistency of gene centrality profile of donors under the same phenotypic group. Silhouette score is computed using function `silhouette` in `R` package `cluster 2.1.4` with euclidean distance as the distance metric.

### False discovery rates for edge estimation

We designed a data-driven permutation experiment to study the false discovery rates of network edges. Given a gene expression matrix D1, the genes are randomly split into two disjoint equal-sized sets, S1 and S2. A “null” dataset D2 is created by permuting the cell ordering of genes in S2 while keeping the original cell ordering of genes in S1. This permutation keeps the overall expression and sparsity levels of the genes the same as in D1 and disassociates genes in S2 from genes in S1 in dataset D2. As a result, the true correlation of gene pairs with one gene in S1 and one gene S2 is 0 in the permuted dataset D2. We denote the estimated correlation matrices of gene pairs with one gene in S1 and one gene in S2 computed with datasets D1 and D2 as *C*1 and *C*2. *C*1 and *C*2 are thresholded to construct the co-expression networks using the 95th and 99th percentiles of the absolute correlations in *C*1, resulting in 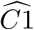 and 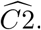. Then, the empirical false discovery rate is computed as

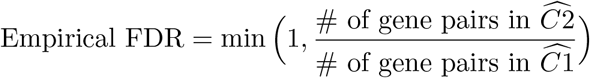

### hTFtarget enrichment

For each transcription factor (TF) present in both the hTFtarget database and the gene co-expression network, we tested the enrichment of its targets from the hTFtarget database (Zhang et al., 2020b) among the genes with edges to the TF in the constructed co-expression network with the standard hypergeometric test. More specifically, for TF *j* with the number of edges *k*_*j*_ in the co-expression network, the number of known targets *m*_*j*_ in the network, the number of correctly identified targets *q*_*j*_, and the total number of genes *n* in the network, the p-value is defined as `phyper`(*q*_*j*_ − 1, *m*_*j*_, *n* − *m*_*j*_, *k*_*j*_, `lower.tail=F`) with the `R` function `phyper`. The hypergeometric test was performed separately for each TF and each donor. The p-values for a TF were combined with the Fisher’s combined probability test (Fisher’s method) as the final p-value of enrichment for the TF.

### PPI enrichment with the STRING database

Protein-protein interaction (PPI) enrichments of the networks were assessed using the proportion of network edges in the STRING database, i.e., the number of edges both in the co-expression network and in the STRING database divided by the number of edges in the co-expression network. A baseline enrichment proportion was computed as

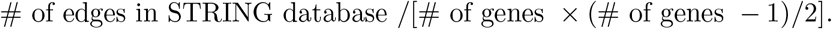

### Differential feature analysis

We performed differential feature analysis on the basis of two types of features: gene expression and gene centrality measures in co-expression networks, including degree, pagerank, betweenness, and eigenvector centrality. Unless otherwise stated, the differential expression tests were implemented at the single cell level using the SCTransform normalized data together with `FindMarkers` function from the `R` package `Seurat` with the `MAST` (Finak et al., 2015) algorithm. Differential centrality tests were carried out at the donor level using the `R` package `Limma` (Smyth, 2005) to accommodate small sample sizes. Donor features “Age” and “Sex” were controlled for in the differential centrality test for the dataset Morabito_2021. Multiplicity correction was performed with the Benjamini-Hochberg procedure (Benjamini and Hochberg, 1995) at a false discovery rate of 0.05 unless otherwise specified.

To investigate the potential confounding of differential network centrality with the differential expression in samples sequenced in two separate batches, we employed DESeq2 (Love et al., 2014) to detect differential expression. Specifically, we implemented DESeq2 through the `FindMarkers` function in Seurat, specifying `test.use` as ”DESeq2”.

### Evaluating the co-expression of Dozer-A3 module genes in immune oligodendrocytes (ODC)

To validate the findings about the Dozer-A3 module of the Morabito_2021, we assessed whether there was higher co-expression of genes in the Dozer-A3 module among immune ODC cells, and whether this association was more significant in the Alzheimer’s group than in the Control group. To address the former, for a given donor *d*, we estimated the probability of a donor ODC cell expressing more than half of the genes in the Dozer-A3 module (*p*_*d*_). We then counted the number of immune ODC cells from that donor (represented by *m*_*d*_) and the number of immune ODC cells expressing at least half of the genes in the Dozer-A3 module (represented by *x*_*d*_). Using these estimates, we assessed the likelihood of observing *x*_*d*_ or more immune ODC cells expressing Dozer-A3 module genes purely by chance, with the following Binomial calculation:

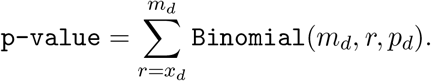

In addition, for a given donor *d*, the association between co-expressing Dozer-A3 module genes and being an immune ODC cell was assessed with a 2 × 2 contingency table. Specifically, a Fisher’s exact test on the contingency table was carried out. The testing of the differences in odds ratios and negative log base 10 transformed Fisher’s exact test p-values of the AD and Control diagnoses were carried out with a Wilcoxon rank sum test.

### Implementation details of the network construction methods

For SCT.Pearson, SCT.Spearman, and Noise.Reg, the gene expression matrix of donor cells were normalized with `SCTransform`. Co-expression matrix for SCT.Pearson (SCT.Spearman) was computed through the Pearson (Spearman) correlation of the normalized counts in the output data slot `scale.data`. The normalized expression in slot `data` was used for Noise.Reg as it requires positive normalized expression. For Noise.Reg, a uniformly distributed random variable from the interval [0, *q*_*i*_), where *q*_*i*_ is the 1st percentile of the normalized expression of gene *i*, was added to the normalized expression before computing Spearman correlation. The normalization for SAVER was performed using the function `saver` and co-expression matrix was obtained with the `saver` function `cor.genes`. Co-expression network for bigSCale2 was computed with function `compute.network`. For MetaCell, gene expression was first normalized with `Seurat`, then projected onto a low-dimensional space with UMAP. Cells were grouped into 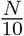 clusters, where *N* is the total number of cells, with the k-means algorithm using the UMAP coordinates. Expression of genes across cells in the same cluster were then aggregated into a meta cell.

The following versions of the software were used for the analysis: `R 4.1.1, SCTransform 0.3.2, SAVER 1.1.2, Seurat 4.0.3, limma 3.48.3, MAST 1.18.0, WGCNA 1.70-3, bigSCale 2.0`.

### Datasets

Jerber_2021. scRNA-seq dataset from Jerber et al. (2021) harbored iPSC cells from multiple donors under neuronal differentiation. Cells were profiled on days 11, 30, and 52 and a phenotypic trait named differentiation efficiency score was derived for each donor. Donors were further classified into two phenotype groups as “success” and “failure” in neuronal differentiation using the differentiation efficiency score. All the personalized gene co-expression network analyses utilized the P_FPP (Proliferating Floor Plate Progenitors) cells on day 11. Donors with fewer than 500 cells, with missing data in differentiation efficiency score, and six donors with extremely low sequencing depths (less than 14% of the median sequencing depth among all donors) were filtered out, leaving a total of 62 donors for the analysis. Genes with an average noise ratio among donors larger than 0.9 were filtered out, resulting in 912 genes per network.

We employed the full Jerber_2021 dataset for personalized co-expression network analysis, and utilized two subsets of the dataset for false discovery analysis and investigating the robustness against sequencing depth differences. The first subset comprised of cells with a depth range between 12k to 17k total UMI counts from 20 donors, while the second subset consisted of 1,139 and 963 cells from a single donor, sequenced in batches labeled “pool2” and “pool3”, respectively.

Morabito 2021. snRNA-seq data of Morabito et al. (2021) were from postmortem human tissues of 11 subjects with Alzheimer disease and and 7 age-matched control subjects. Oligodendrocytes, which accounted for 60% of the total number of cells in the datasets (with a median number of 2,005 cells per donor) were utilized in all the the co-expression analyses. To facilitate a direct comparison with the co-expression network analysis conducted in the data paper (scWGCNA), the same set of 1,252 genes were used as the starting gene set and were further filtered out if they satisfied any of the following conditions: (i) expressed in less than 2 cells in any donor; (ii) average noise ratio over donors is larger than 0.85. This resulted in 570 genes per donor for the analysis.

Cuomo 2020. scRNA-seq data of iPSC cells from Cuomo et al. (2020) were used to generate simulation parameters. Cells in this dataset have high sequencing depths (averaging approximately 530K total counts per cell), allowing estimation of more realistic parameters to serve as ground truth in simulations.

## Data access

Single cell RNA-Seq dataset Jerber_2021 (Jerber et al., 2021) is available at https://zenodo.org/record/4333872#.YnhH6HXMJH4, dataset Cuomo_2020 is available at https://zenodo.org/record/3625024#.YnhMfnXMJH4, and dataset `Morabito_202` is available at https://www.synapse.org/#!Synapse:syn22079621/.

Dozer `R` package and vignette are available at https://github.com/keleslab/Dozer.

## Acknowledgements

This work was supported by National Institutes of Health (NIH) grant HG003747 and a Chan Zuckerberg Initiative (CZI) Single Cell Data Insights grant.

## References

Arrázola MS, Andraini T, Szelechowski M, Mouledous L, Arnauné-Pelloquin L, Davezac N, Belenguer P, Rampon C, and Miquel MC. 2019. Mitochondria in developmental and adult neurogenesis. Neurotoxicity Research 36: 257–267.

Azevedo C, Teku G, Pomeshchik Y, Reyes JF, Chumarina M, Russ K, Savchenko E, Hammarberg A, Lamas NJ, Collin A, et al. 2022. Parkinson’s disease and multiple system atrophy patient iPSC-derived oligodendrocytes exhibit alpha-synuclein–induced changes in maturation and immune reactive properties. Proceedings of the National Academy of Sciences 119: e2111405119.

Azuaje FJ. 2014. Selecting biologically informative genes in co-expression networks with a centrality score. Biology direct 9: 1–23.

Bacher R, Chu LF, Leng N, Gasch AP, Thomson JA, Stewart RM, Newton M, and Kendziorski C. 2017. SCnorm: robust normalization of single-cell RNA-seq data. Nature methods 14: 584–586.

Ball Jr AR, Chen YY, and Yokomori K. 2014. Mechanisms of cohesin-mediated gene regulation and lessons learned from cohesinopathies. Biochimica et Biophysica Acta (BBA)-Gene Regulatory Mechanisms 1839: 191–202.

Baran Y, Bercovich A, Sebe-Pedros A, Lubling Y, Giladi A, Chomsky E, Meir Z, Hoichman M, Lifshitz A, and Tanay A. 2019. MetaCell: analysis of single-cell RNA-seq data using K-nn graph partitions. Genome biology 20: 1–19.

Becht E, McInnes L, Healy J, Dutertre CA, Kwok IW, Ng LG, Ginhoux F, and Newell EW. 2019. Dimensionality reduction for visualizing single-cell data using UMAP. Nature biotechnology 37: 38–44.

Benjamini Y and Hochberg Y. 1995. Controlling The False Discovery Rate - A Practical And Powerful Approach To Multiple Testing. J. Royal Statist. Soc., Series B 57: 289 –300.

Bernardes JP, Mishra N, Tran F, Bahmer T, Best L, Blase JI, Bordoni D, Franzenburg J, Geisen U, Josephs-Spaulding J, et al. 2020. Longitudinal Multi-omics Analyses Identify Responses of Megakaryocytes, Erythroid Cells, and Plasmablasts as Hallmarks of Severe COVID-19. Immunity 53: 1296–1314.e9.

Brunetti D, Dykstra W, Le S, Zink A, and Prigione A. 2021. Mitochondria in neurogenesis: Implications for mitochondrial diseases. Stem Cells 39: 1289–1297.

Chen YJJ, Friedman BA, Ha C, Durinck S, Liu J, Rubenstein JL, Seshagiri S, and Modrusan Z. 2017. Single-cell RNA sequencing identifies distinct mouse medial ganglionic eminence cell types. Scientific reports 7: 1–11.

Choudhary S and Satija R. 2022. Comparison and evaluation of statistical error models for scRNA-seq. Genome biology 23.

Cuomo AS, Seaton DD, McCarthy DJ, Martinez I, Bonder MJ, Garcia-Bernardo J, Amatya S, Madrigal P, Isaacson A, Buettner F, et al. 2020. Single-cell RNA-sequencing of differentiating iPS cells reveals dynamic genetic effects on gene expression. Nature communications 11: 1–14.

Eraslan G, Simon LM, Mircea M, Mueller NS, and Theis FJ. 2019. Single-cell RNA-seq denoising using a deep count autoencoder. Nature communications 10: 1–14.

Finak G, McDavid A, Yajima M, Deng J, Gersuk V, Shalek AK, Slichter CK, Miller HW, McElrath MJ, Prlic M, et al. 2015. MAST: a flexible statistical framework for assessing transcriptional changes and characterizing heterogeneity in single-cell RNA sequencing data. Genome biology 16: 1–13.

Forbes AN. 2022. Discovery of novel therapeutic targets in cancer using patient-specific gene regulatory networks. Ph.D. thesis, Weill Medical College of Cornell University.

Fujita Y, Masuda K, Bando M, Nakato R, Katou Y, Tanaka T, Nakayama M, Takao K, Miyakawa T, Tanaka T, et al. 2017. Decreased cohesin in the brain leads to defective synapse development and anxiety-related behavior. Journal of Experimental Medicine 214: 1431–1452.

Guo C, Sun L, Chen X, and Zhang D. 2013. Oxidative stress, mitochondrial damage and neurodegenerative diseases. Neural regeneration research 8: 2003.

Hafemeister C and Satija R. 2019. Normalization and variance stabilization of single-cell RNA-seq data using regularized negative binomial regression. Genome biology 20: 1–15.

He X and Zhang J. 2006. Why do hubs tend to be essential in protein networks? PLoS genetics 2: e88.

Hroudová J, Singh N, Fišar Z, et al. 2014. Mitochondrial dysfunctions in neurodegenerative diseases: relevance to Alzheimer’s disease. BioMed research international 2014.

Huang M, Wang J, Torre E, Dueck H, Shaffer S, Bonasio R, Murray JI, Raj A, Li M, and Zhang NR. 2018. SAVER: gene expression recovery for single-cell RNA sequencing. Nature methods 15: 539–542.

Iacono G, Massoni-Badosa R, and Heyn H. 2019. Single-cell transcriptomics unveils gene regulatory network plasticity. Genome biology 20: 1–20.

Jeong H, Mason SP, Barabási AL, and Oltvai ZN. 2001. Lethality and centrality in protein networks. Nature 411: 41–42.

Jerber J, Seaton DD, Cuomo AS, Kumasaka N, Haldane J, Steer J, Patel M, Pearce D, Andersson M, Bonder MJ, et al. 2021. Population-scale single-cell RNA-seq profiling across dopaminergic neuron differentiation. Nature genetics 53: 304–312.

Johnson KA and Krishnan A. 2022. Robust normalization and transformation techniques for constructing gene coexpression networks from RNA-seq data. Genome biology 23: 1–26.

Kausar S, Wang F, and Cui H. 2018. The role of mitochondria in reactive oxygen species generation and its implications for neurodegenerative diseases. Cells 7: 274.

Kirby L, Jin J, Cardona JG, Smith MD, Martin KA, Wang J, Strasburger H, Herbst L, Alexis M, Karnell J, et al. 2019. Oligodendrocyte precursor cells present antigen and are cytotoxic targets in inflammatory demyelination. Nature communications 10: 1–20.

Kuleshov MV, Jones MR, Rouillard AD, Fernandez NF, Duan Q, Wang Z, Koplev S, Jenkins SL, Jagodnik KM, Lachmann A, et al. 2016. Enrichr: a comprehensive gene set enrichment analysis web server 2016 update. Nucleic acids research 44: W90–W97.

Langfelder P and Horvath S. 2008. WGCNA: an R package for weighted correlation network analysis. BMC bioinformatics 9: 1–13.

Lareau CA, White BC, Oberg AL, and McKinney BA. 2015. Differential co-expression network centrality and machine learning feature selection for identifying susceptibility hubs in networks with scale-free structure. BioData mining 8: 1–17.

Liu S, Liu Y, Hao W, Wolf L, Kiliaan AJ, Penke B, Rübe CE, Walter J, Heneka MT, Hartmann T, et al. 2012. TLR2 is a primary receptor for Alzheimer’s amyloid β peptide to trigger neuroinflammatory activation. The Journal of Immunology 188: 1098–1107.

Love MI, Huber W, and Anders S. 2014. Moderated estimation of fold change and dispersion for RNA-seq data with DESeq2. Genome biology 15: 1–21.

Mahajani S, Giacomini C, Marinaro F, De Pietri Tonelli D, Contestabile A, and Gasparini L. 2017. Lamin B1 levels modulate differentiation into neurons during embryonic corticogenesis. Scientific reports 7: 1–11.

Morabito S, Miyoshi E, Michael N, Shahin S, Martini AC, Head E, Silva J, Leavy K, Perez-Rosendahl M, and Swarup V. 2021. Single-nucleus chromatin accessibility and transcriptomic characterization of Alzheimer’s disease. Nature Genetics 53: 1143–1155.

Mostafavi S, Gaiteri C, Sullivan SE, White CC, Tasaki S, Xu J, Taga M, Klein HU, Patrick E, Komashko V, et al. 2018. A molecular network of the aging human brain provides insights into the pathology and cognitive decline of Alzheimer’s disease. Nature neuroscience 21: 811–819.

Nagy C, Maitra M, Tanti A, Suderman M, Théroux JF, Davoli MA, Perlman K, Yerko V, Wang YC, Tripathy SJ, et al. 2020. Single-nucleus transcriptomics of the prefrontal cortex in major depressive disorder implicates oligodendrocyte precursor cells and excitatory neurons. Nature neuroscience 23: 771–781.

Pratapa A, Jalihal AP, Law JN, Bharadwaj A, and Murali T. 2020. Benchmarking algorithms for gene regulatory network inference from single-cell transcriptomic data. Nature methods 17: 147–154.

Risso D, Perraudeau F, Gribkova S, Dudoit S, and Vert JP. 2018. A general and flexible method for signal extraction from single-cell RNA-seq data. Nature communications 9: 1–17.

Robinson MD and Oshlack A. 2010. A scaling normalization method for differential expression analysis of RNA-seq data. Genome biology 11: 1–9.

Rousseeuw PJ. 1987. Silhouettes: a graphical aid to the interpretation and validation of cluster analysis. Journal of computational and applied mathematics 20: 53–65.

Sarkar A and Stephens M. 2021. Separating measurement and expression models clarifies confusion in single-cell RNA sequencing analysis. Nature genetics 53: 770–777.

Savino A, Provero P, and Poli V. 2020. Differential co-expression analyses allow the identification of critical signalling pathways altered during tumour transformation and progression. International journal of molecular sciences 21: 9461.

Shi Y and Holtzman DM. 2018. Interplay between innate immunity and Alzheimer disease: APOE and TREM2 in the spotlight. Nature Reviews Immunology 18: 759–772.

Smyth GK. 2005. Limma: linear models for microarray data. In Bioinformatics and computational biology solutions using R and Bioconductor, pp. 397–420. Springer.

Soskic B, Cano-Gamez E, Smyth DJ, Ambridge K, Ke Z, Matte JC, Bossini-Castillo L, Kaplanis J, Ramirez-Navarro L, Lorenc A, et al. 2022. Immune disease risk variants regulate gene expression dynamics during CD4+ T cell activation. Nature Genetics 54: 817–826.

Stone M, McCalla SG, Siahpirani AF, Periyasamy V, Shin J, and Roy S. 2021. Identifying strengths and weaknesses of methods for computational network inference from single cell RNA-seq data. bioRxiv.

Sun T, Song D, Li WV, and Li JJ. 2021. scDesign2: a transparent simulator that generates high-fidelity single-cell gene expression count data with gene correlations captured. Genome biology 22: 163.

Szklarczyk D, Gable AL, Nastou KC, Lyon D, Kirsch R, Pyysalo S, Doncheva NT, Legeay M, Fang T, Bork P, et al. 2021. The STRING database in 2021: customizable protein–protein networks, and functional characterization of user-uploaded gene/measurement sets. Nucleic acids research 49: D605–D612.

Tran HTN, Ang KS, Chevrier M, Zhang X, Lee NYS, Goh M, and Chen J. 2020. A benchmark of batch-effect correction methods for single-cell RNA sequencing data. Genome biology 21: 1–32.

Travaglini KJ, Nabhan AN, Penland L, Sinha R, Gillich A, Sit RV, Chang S, Conley SD, Mori Y, Seita J, et al. 2020. A molecular cell atlas of the human lung from single-cell RNA sequencing. Nature 587: 619–625.

Vallejos CA, Risso D, Scialdone A, Dudoit S, and Marioni JC. 2017. Normalizing single-cell RNA sequencing data: challenges and opportunities. Nature methods 14: 565–571.

Van Der Wijst MG, Brugge H, De Vries DH, Deelen P, Swertz MA, and Franke L. 2018. Single-cell RNA sequencing identifies cell type-specific cis-eQTLs and co-expression QTLs. Nature genetics 50: 493–497.

Van Dijk D, Sharma R, Nainys J, Yim K, Kathail P, Carr AJ, Burdziak C, Moon KR, Chaffer CL, Pattabiraman D, et al. 2018. Recovering gene interactions from single-cell data using data diffusion. Cell 174: 716–729.

Wang J, Huang M, Torre E, Dueck H, Shaffer S, Murray J, Raj A, Li M, and Zhang NR. 2018. Gene expression distribution deconvolution in single-cell RNA sequencing. Proceedings of the National Academy of Sciences 115: E6437–E6446.

Wang X, Choi D, and Roeder K. 2021. Constructing local cell-specific networks from single-cell data. Proceedings of the National Academy of Sciences 118.

van der Wijst MG, de Vries DH, Groot HE, Trynka G, Hon CC, Bonder MJ, Stegle O, Nawijn M, Idaghdour Y, van der Harst P, et al. 2020. Science Forum: The single-cell eQTLGen consortium. Elife 9: e52155.

Zeisel A, Muñoz-Manchado AB, Codeluppi S, L önnerberg P, La Manno G, Juréus A, Marques S, Munguba H, He L, Betsholtz C, et al. 2015. Cell types in the mouse cortex and hippocampus revealed by single-cell RNA-seq. Science 347: 1138–1142.

Zhang B and Horvath S. 2005. A general framework for weighted gene co-expression network analysis. Statistical applications in genetics and molecular biology 4.

Zhang MJ, Ntranos V, and Tse D. 2020a. Determining sequencing depth in a single-cell RNA-seq experiment. Nature communications 11: 1–11.

Zhang Q, Liu W, Zhang HM, Xie GY, Miao YR, Xia M, and Guo AY. 2020b. hTFtarget: a comprehensive database for regulations of human transcription factors and their targets. Genomics, proteomics & bioinformatics 18: 120–128.

Zhang R, Atwal GS, and Lim WK. 2021. Noise regularization removes correlation artifacts in single-cell RNA-seq data preprocessing. Patterns 2: 100211.

Zheng S, Cheng G, Guo J, and Zhu H. 2019. Test for high dimensional correlation matrices. Annals of statistics 47: 2887.

