## Supplementary materials for "Dozer: Debiased personalized gene co-expression networks for population-scale scRNA-seq data"

Shan Lu[1] and Sündüz Keleş[1,2]

[1] Department of Statistics, University of Wisconsin, Madison, WI, USA.

[2] Department of Biostatistics and Medical Informatics, University of Wisconsin School of Medicine and Public Health, Madison, WI, USA.

#### Contents

**1 Results for computational experiments** **2**

**2 Additional results for the analysis of the `Jerber_2021` dataset** **13**

**3 Additional results for the analysis of the `Morabito_2021` dataset** **17**

### 1 Results for computational experiments

#### 1.1 Analytical form of the gene noise ratio and correction factor under the Gamma-Poisson model

Under the Gamma-Poisson model, the noise ratio $R_j$ and correction factor $S_j$, $j = 1, \cdots, G$ can be expressed as functions of the cell sequencing depths $\{\ell_i\}_{i=1}^N$ and the Gamma scale parameter $u_j$ as:

$$R_j = \frac{1}{1 + \frac{u_j}{\frac{1}{N}\sum_{i=1}^N 1/\ell_i}}, \tag{1}$$

and

$$S_j = 1 + \frac{\sum_{i=1}^N 1/\ell_i}{N} \times \frac{1}{u_j}. \tag{2}$$

These analytical forms in equations (1) and (2) support the intuition and the empirical results that when the sequencing depth or expression is high, $R_j$ is closer to 0 ($S_j$ is closer to 1) and the bias of sample correlation of normalized UMI counts is expected to be small. Figure S1 displays the noise ratio as a function of gene expression and sparsity (i.e., proportion of zero counts for a gene) across both simulated and real datasets (`Jerber_2021` and `Morabito_2021`) and confirms the expected pattern of noise ratio as mean expression level and sparsity vary. We observe that for a fixed expression level (x-axis), genes with lower sparsity tend to have higher noise ratios. While this initially appears counter-intuitive, further calculations with this model support the observation. Under the Gamma-Poisson model with $v_j$, $u_j$ and $\ell_i$ as gamma shape, scale parameters for gene $j$ and sequencing depth for cell $i$, the probability of zero count for gene $j$ is

$$\mathbb{P}(Y_{ij} = 0) = \left(\frac{1}{1 + \ell_i u_j}\right)^{v_j}. \tag{3}$$

For a fixed mean expression $c_0$, i.e., $u_j v_j = c_0$,

$$\mathbb{P}(Y_{ij} = 0) = \frac{1}{\left(1 + \frac{c_0 \ell_i}{v_j}\right)^{v_j}}. \tag{4}$$

The denominator of eqn. (4) is a monotonically increasing function of $v_j$ for fixed $c_0\ell_i$. As a result, $\mathbb{P}(Y_{ij} = 0)$ decreases as $v_j$ increases and, for fixed mean expression $c_0 = u_j v_j$, the sparsity level of gene $j$ increases as $u_j$ increases. Further utilizing eqn. (1), we conclude that $R_j$ decreases as $u_j$ increases; hence, for fixed mean expression, sparser genes have lower noise ratio, which matches the empirical observation in Figure S1.

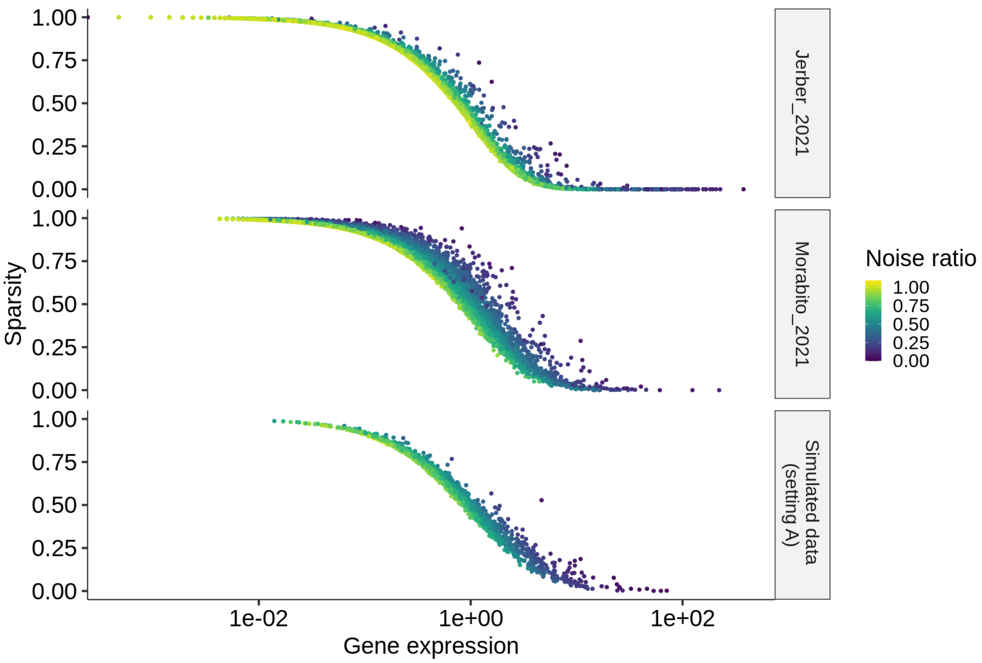


Figure S1: Noise ratio as a function of mean gene expression, sparsity level (average zero count proportion) across all genes without filtering for high noise ratio in the `Jerber_2021`, `Morabito_2021`, and simulated datasets.

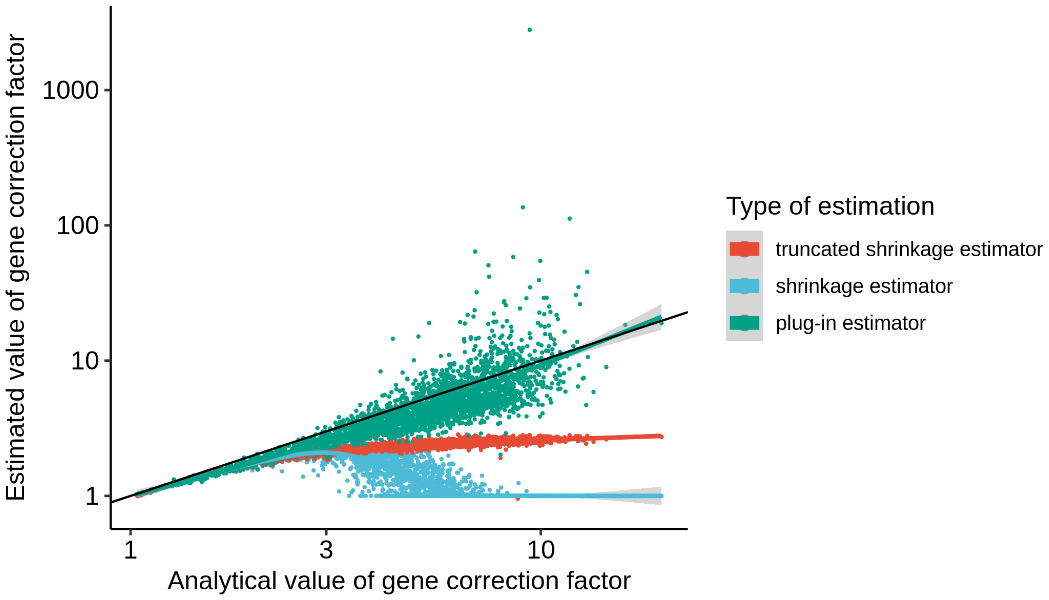


Figure S2: Comparison of the analytical and estimated correction factors $S_j$, $j = 1, \cdots, G$ in the simulation study from the Gamma-Poisson base model. Three different estimators of the correction factors are depicted by different colors. The initial plug-in estimator (Eqn. 14, green) is noisy for large values of the correction factor. The shrinkage estimator (Eqn. 16, blue) steers the large scale factors down to 1. The truncated shrinkage estimator (Eqn. 17, red) reduces the variation and keeps the monotonicity of the estimate.

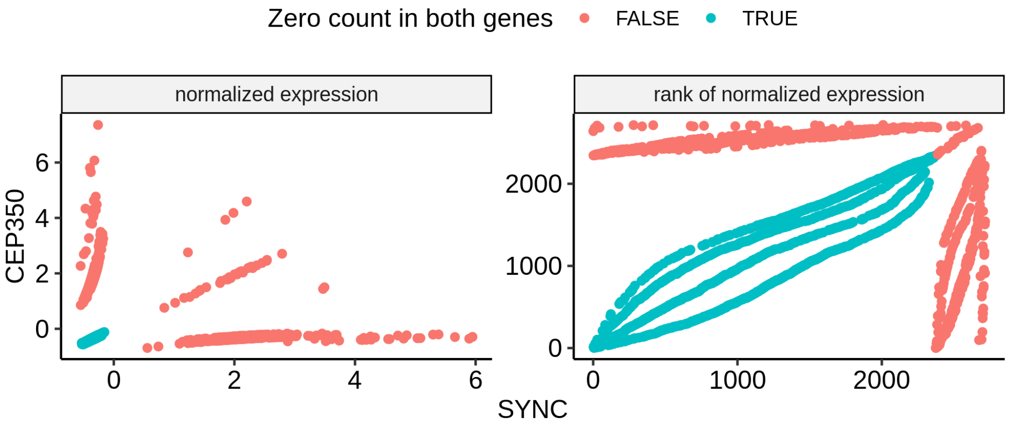


Figure S3: Illustrative over-smoothing resulting from the Spearman correlation computed with the SCTransform normalized counts of P_FPP cells in the `Jerber_2021` dataset. Genes *SYNC* and *CEP350* both have sparse UMI counts with zero expression in 72% and 80% of the cells, respectively. Left and right panels display the scatter plots of SCTranform normalized expression and the corresponding ranks of the normalized expression of the two genes, respectively. The Pearson and Spearman correlations of the two genes are -0.03 and 0.46, respectively. Cells with zero expression for both genes are depicted in blue and lead to positive Spearman correlation for these two genes. This is in contradiction with the Chi-square test which tests the independence of zero or positive expression between the two genes and results in a p-value of 0.49.

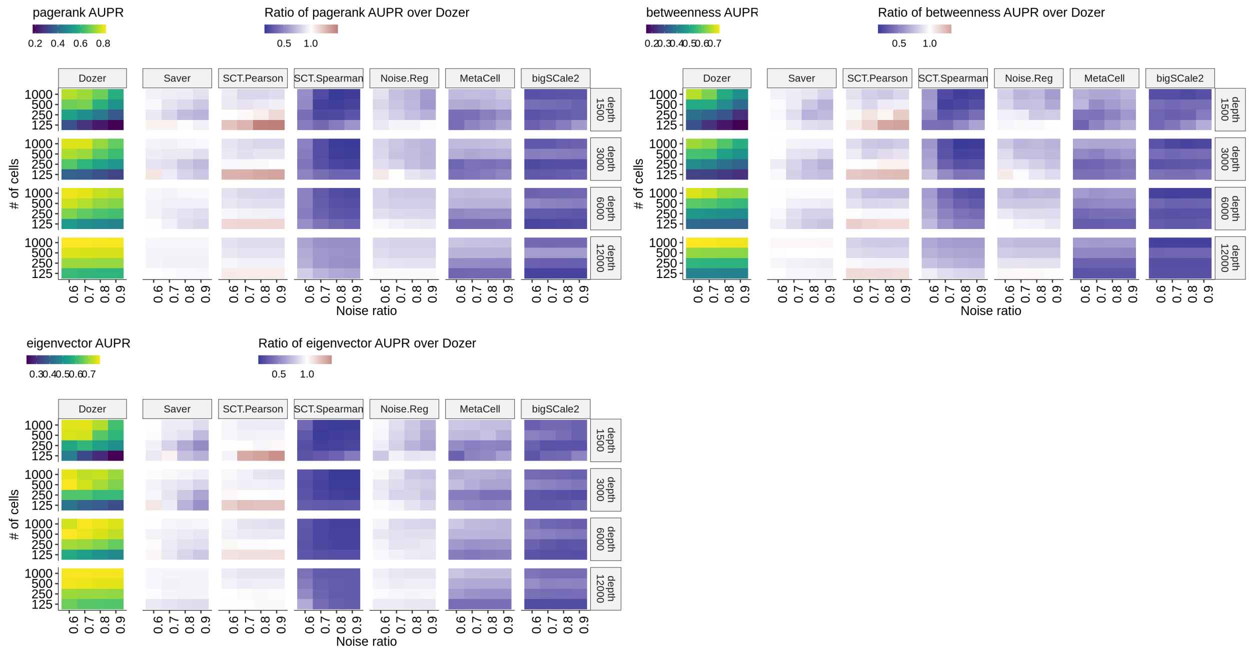


Figure S4: Summary of simulation setting A results in terms of AUPR scores for high centrality gene identification. The average AUPR scores for the identification of genes with top gene centralities as a function of gene noise ratios, cell sample sizes, and average sequencing depths. Gene centrality measures include pagerank, betweenness, and eigenvector centrality. The left most panel depicts the average AUPR score of Dozer. The other panels highlight the performances of other methods as quantified by the ratio of their AUPR scores over the AUPR score of Dozer.

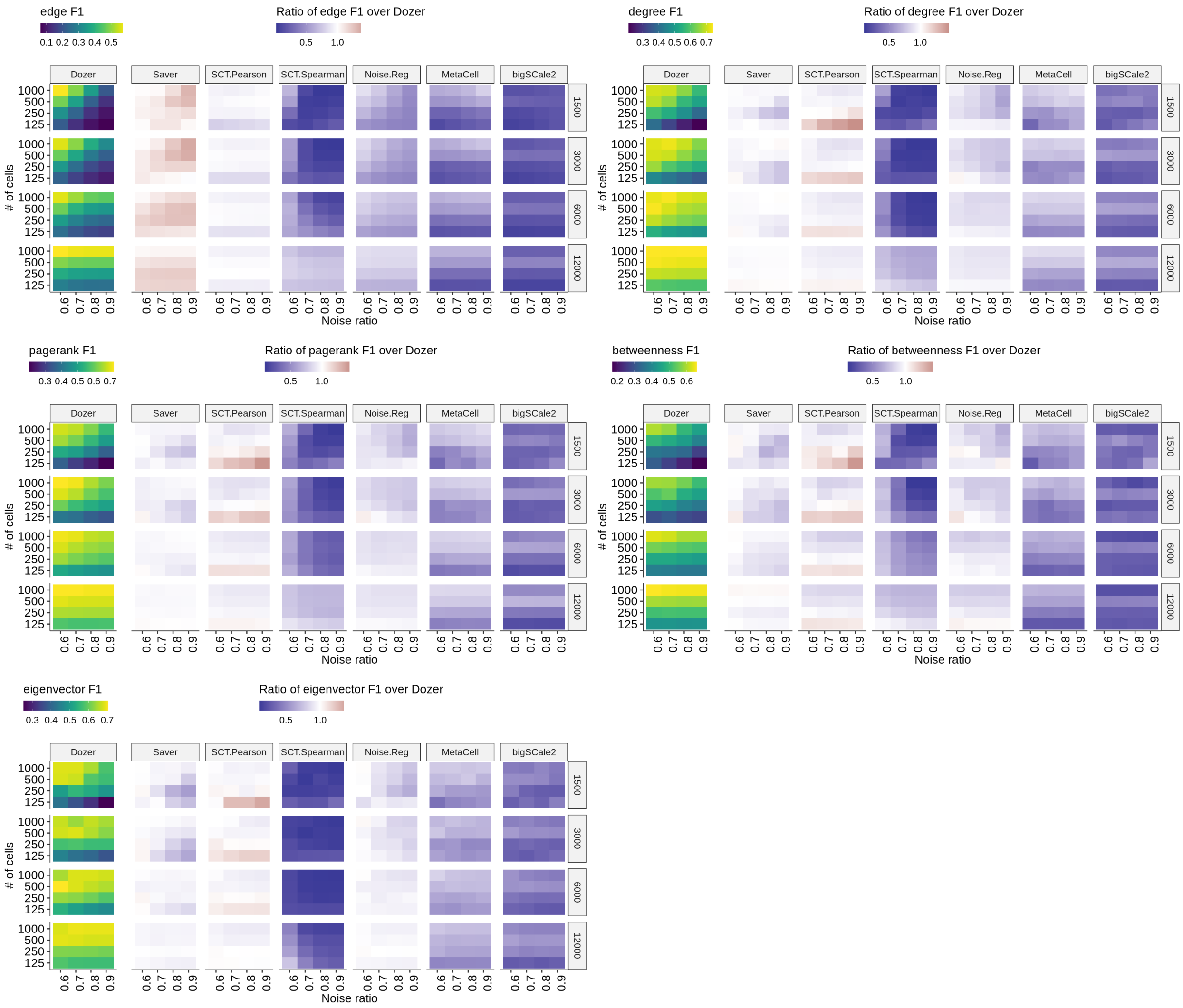


Figure S5: Summary of simulation setting A results in terms of F1 scores for edge and high centrality gene identification. The average F1 scores for edge and high centrality gene identification as a function of gene noise ratios, cell sample sizes and average sequencing depths. Gene centrality measures include degree, pagerank, betweenness, and eigenvector centralities. The left panel shows the average F1 score of Dozer. The other panels highlight the performances of other methods as quantified by the ratio of their F1 scores over the F1 score of Dozer.

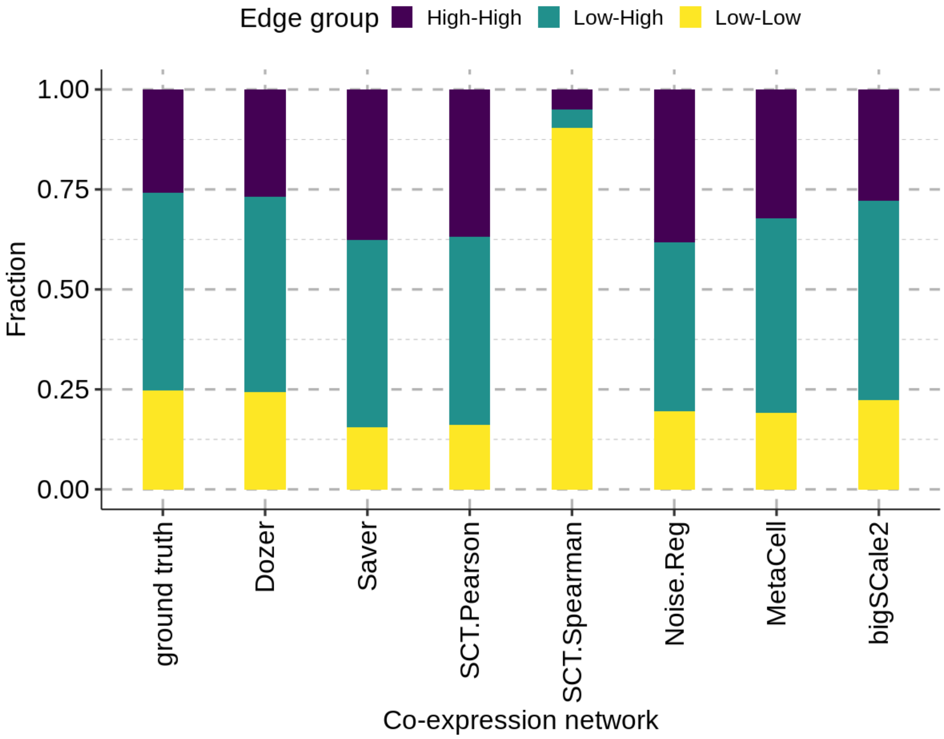


Figure S6: Stratification of network edges inferred by each method with respect to expression groups of the gene pairs: High-High: both genes in high expression group; Low-Low: both genes in low expression group; High-Low: one gene in high and the other in the low expression group. Y-axis depicts the fraction of each method's inferred network edges in each expression group. High (low) expression group is defined as the set of genes with expression larger (smaller) than the median expression among all genes. Data is generated from simulation setting A with true network edges corresponding to the gene pairs with absolute correlation exceeding the 95% percentile in the true correlation matrix in magnitude.

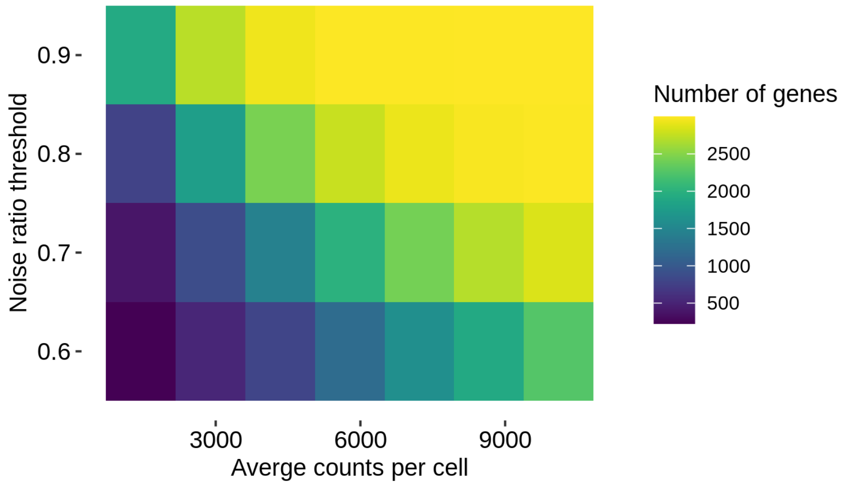


Figure S7: Quantification of numbers of genes with varying levels of noise ratios as a function of average sequencing depth per cell. Colored entries depict numbers of genes with noise ratios smaller than the y-axis value. Quantification is done on data from simulation setting A and includes a total of 3,000 genes.

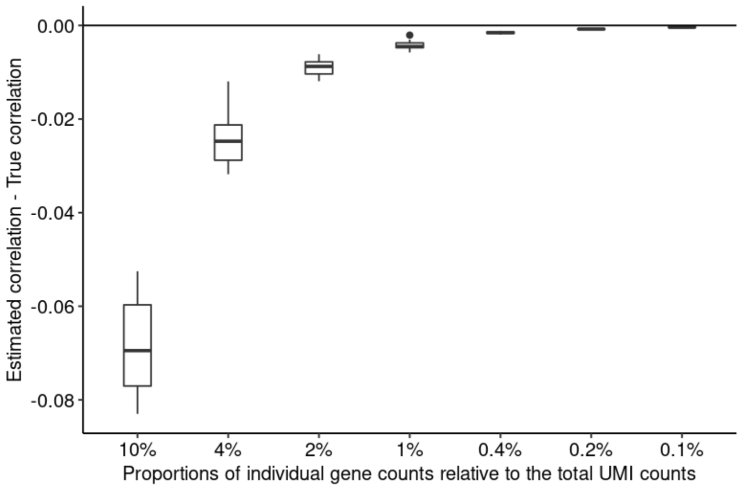


Figure S8: Boxplots for the bias in estimated gene pair correlations induced by the normalization procedure using total UMI counts as a global cell size factor. The data is simulated using the package `Splatter` [3]. The simulation parameters are taken from the example object `newSplatParams` in `Splatter`. In the simulation, the true correlations between gene pairs are set to zero. Gene pair correlations are computed using expression values normalized by total UMI counts. The estimation bias is calculated for seven different gene proportion scenarios (x-axis), ranging from genes accounting for 10% to 0.1% of the total UMI counts.

#### 1.2 Simulation experiment with scDesign2 [2]

To evaluate robustness of Dozer to potential violations from the Poisson-Gamma setting, we leveraged another scRNA-seq simulation tool, `scDesign2` [2], which does not solely rely on the Poisson-Gamma setting. `ScDesign2` chooses the marginal distributions of the genes adaptively among a larger set of count distributions: Poisson-Gamma, Negative Binomial-Gamma, Zero Inflated Poisson-Gamma, and Zero Inflated Negative Binomial and employs a Copula model to generate gene-gene correlations. We used the P_FPP cells of donor "HPSI0114i-eipl_1" in Jerber_2021 dataset to estimate the simulation parameters for `scDesign2`. `scDesign2` parameters were set to their defaults following the `scDesign2` vignette, with the exception of randomizing the gene ordering for marginal distributions. This was done to disentangle the relationships between gene correlations and marginal distributions. This yielded 33, 130, and 2,053 genes with Poisson, Zero inflated Negative Binomial, and Negative Binomial as estimated distributions, respectively. Overall, in this more flexible simulation setting, Dozer performed better than the alternatives in the identification of network edges (except under the low number of cells (less than 250 cells) settings where Saver has a slight advantage) and high centrality genes (Supplementary Figures S9, S10 for AUPR and F1 scores, respectively).

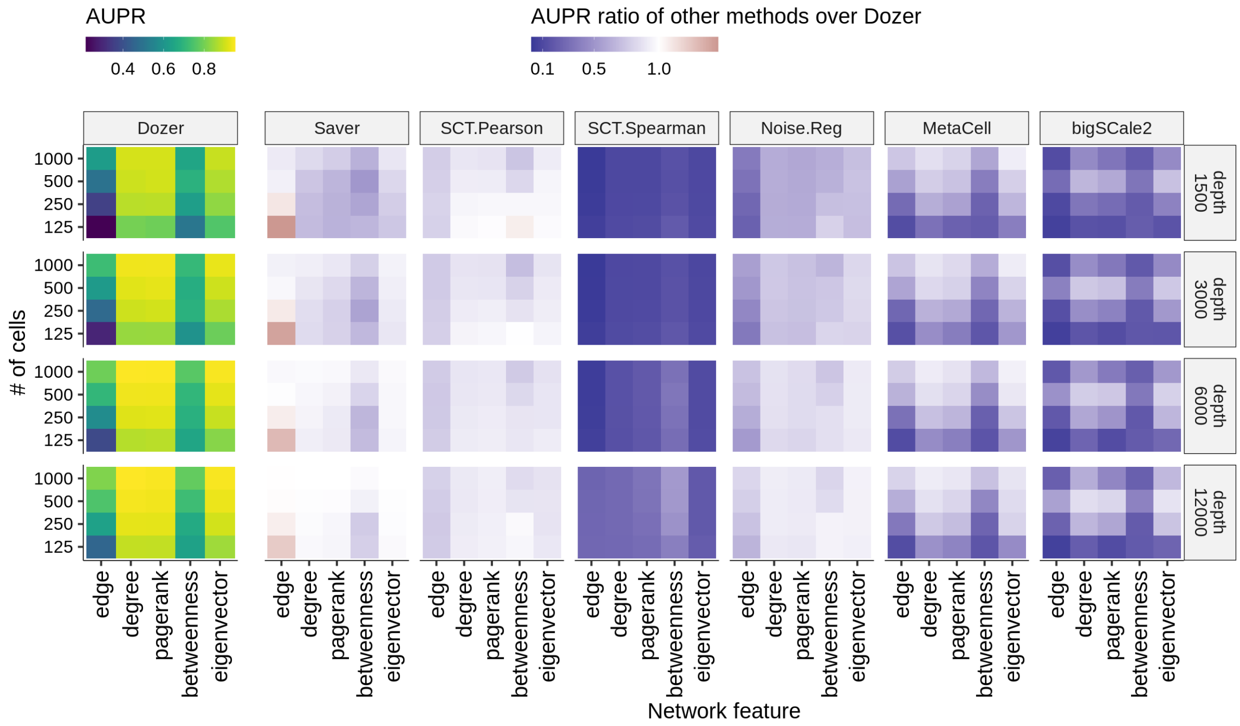


Figure S9: Summary of the scDesign2 simulations in terms of AUPR scores for edge and high centrality gene identification. The average AUPR scores for edge and high centrality gene identification as a function of cell sample sizes and average sequencing depths. Gene centrality measures include degree, pagerank, betweenness, and eigenvector centralities. The left panel shows the average AUPR score of Dozer. The other panels highlight the performances of other methods as quantified by the ratio of their AUPR scores over the AUPR score of Dozer.

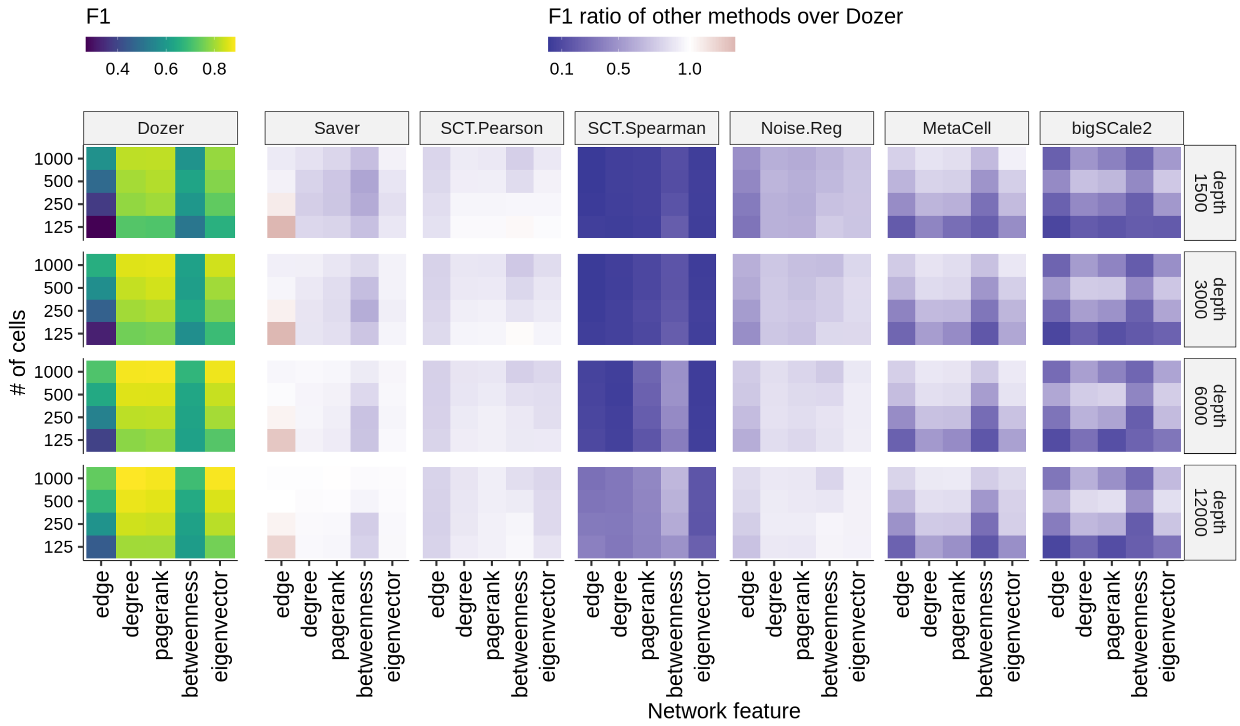


Figure S10: Summary of the `scDesign2` simulations in terms of F1 scores for edge and high centrality gene identification. The average F1 scores for edge and high centrality gene identification as a function of cell sample sizes and average sequencing depths. Gene centrality measures include degree, pagerank, betweenness, and eigenvector centralities. The left panel shows the average F1 score of Dozer. The other panels highlight the performances of other methods as quantified by the ratio of their F1 scores over the F1 score of Dozer.

### 2 Additional results for the analysis of the Jerber_2021 dataset

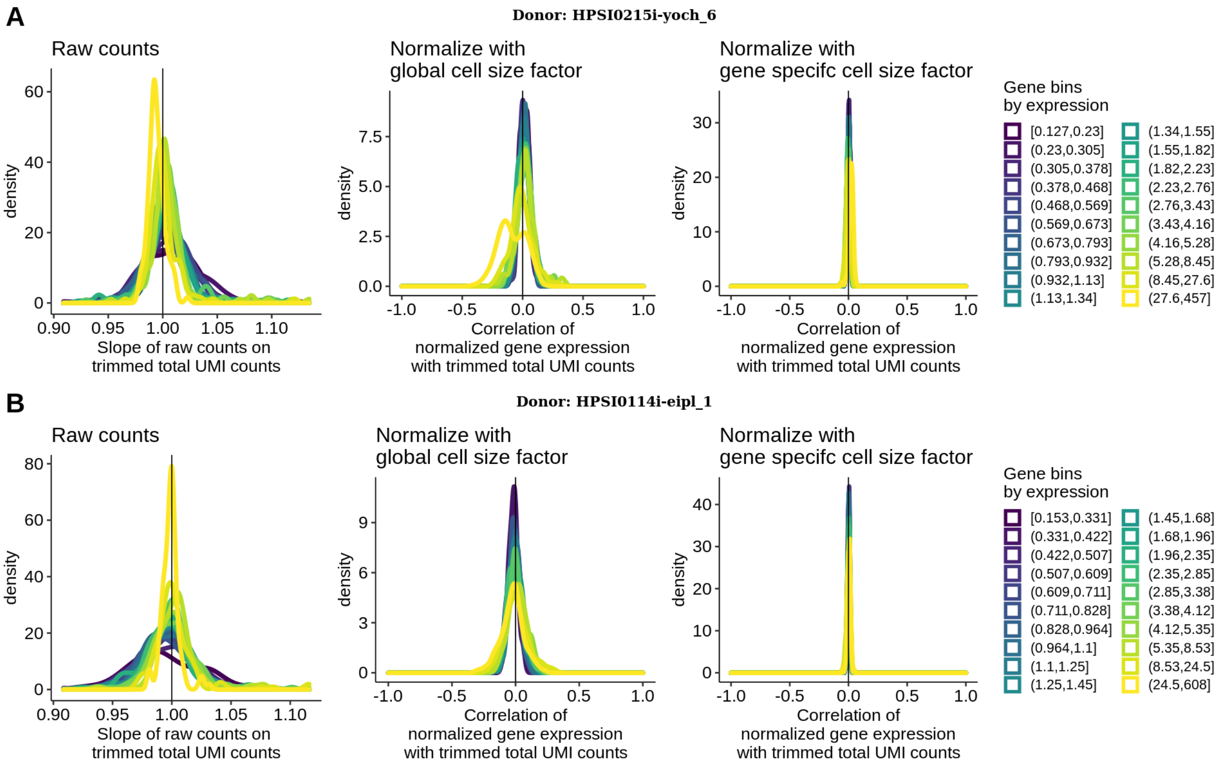


Figure S11: Diagnostics for cell size factor estimation. Panels A and B represent datasets from two different donors. The left most panels of A and B display the density of regression slopes between raw counts and trimmed total UMI counts in gene groups stratified by expression. The middle (right) panels of A and B show the density of correlations between normalized expression with a global cell size factor (with gene specific cell size factors) and trimmed total UMI counts in gene groups stratified by expression.

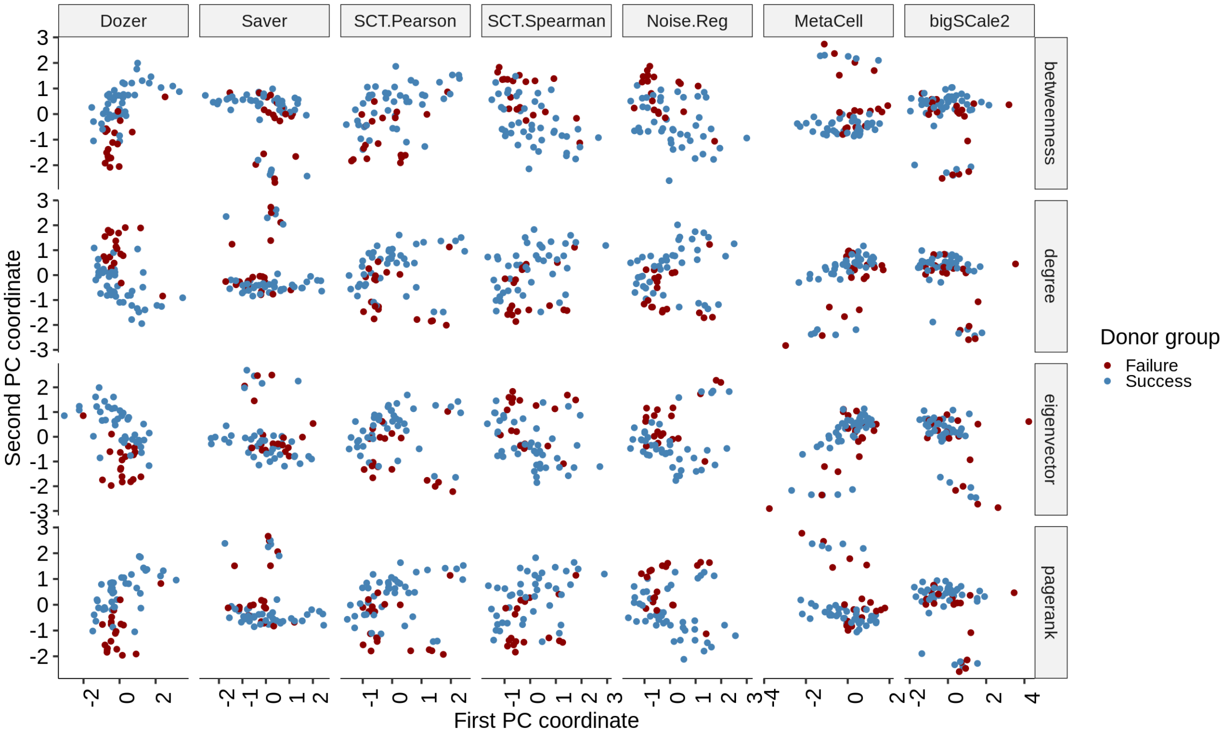


Figure S12: Two-dimensional projection of the `Jerber_2021` donor networks based on their gene centrality measures with principal components (PC). Color of the data points represents phenotypic groups, i.e., failure and success in neuronal differentiation.

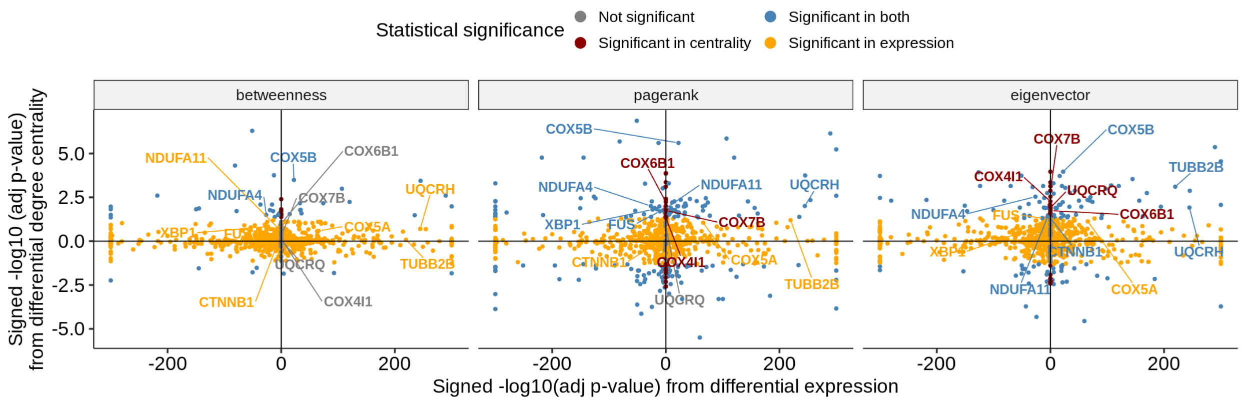


Figure S13: Comparison of differential centralities of the genes from the Dozer co-expression networks with their differential expression. Centrality measures include betweenness, pagerank and eigenvector centrality. X-axis displays the signed -log10(adjusted p-values) from differential expression with positive (negative) values denoting higher (lower) expression in the Failure group. Y-axis denotes the signed -log10(adjusted p-value) from differential degree centrality with positive (negative) values exhibiting higher (lower) centrality in the Failure group. The differential degree genes enriched in the KEGG term "Pathways of neurodegeneration" (Supplementary Figure S14) are labeled in the figure, demonstrating the consistency among various centrality measures.

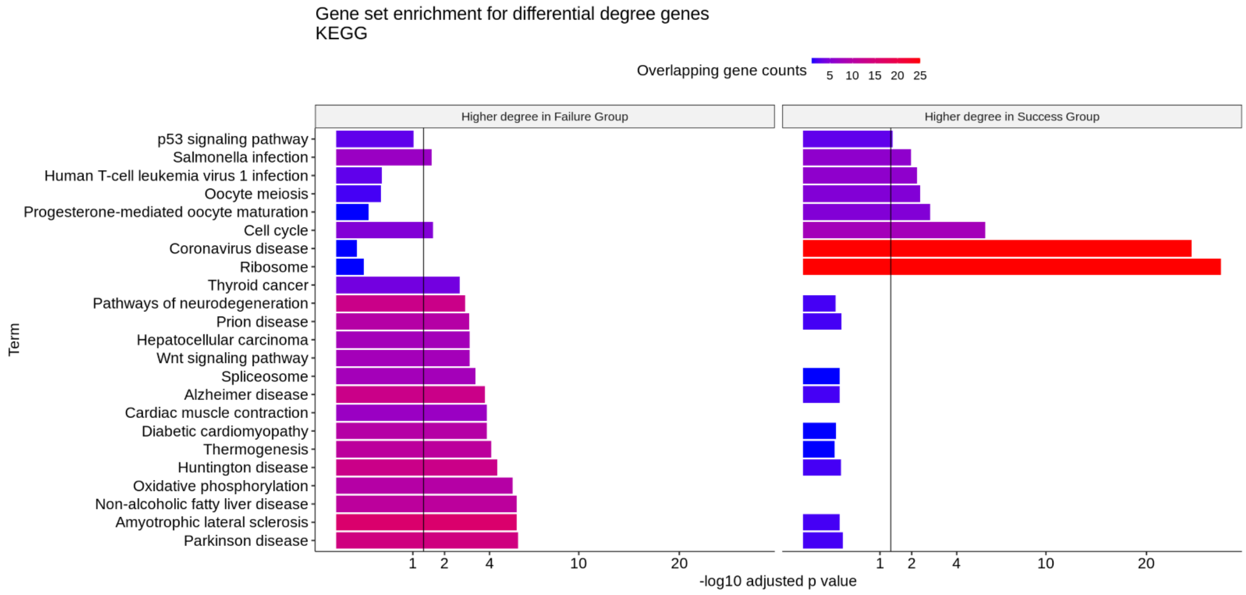


Figure S14: Gene set enrichment analysis of the Dozer identified differential degree genes with KEGG pathways.

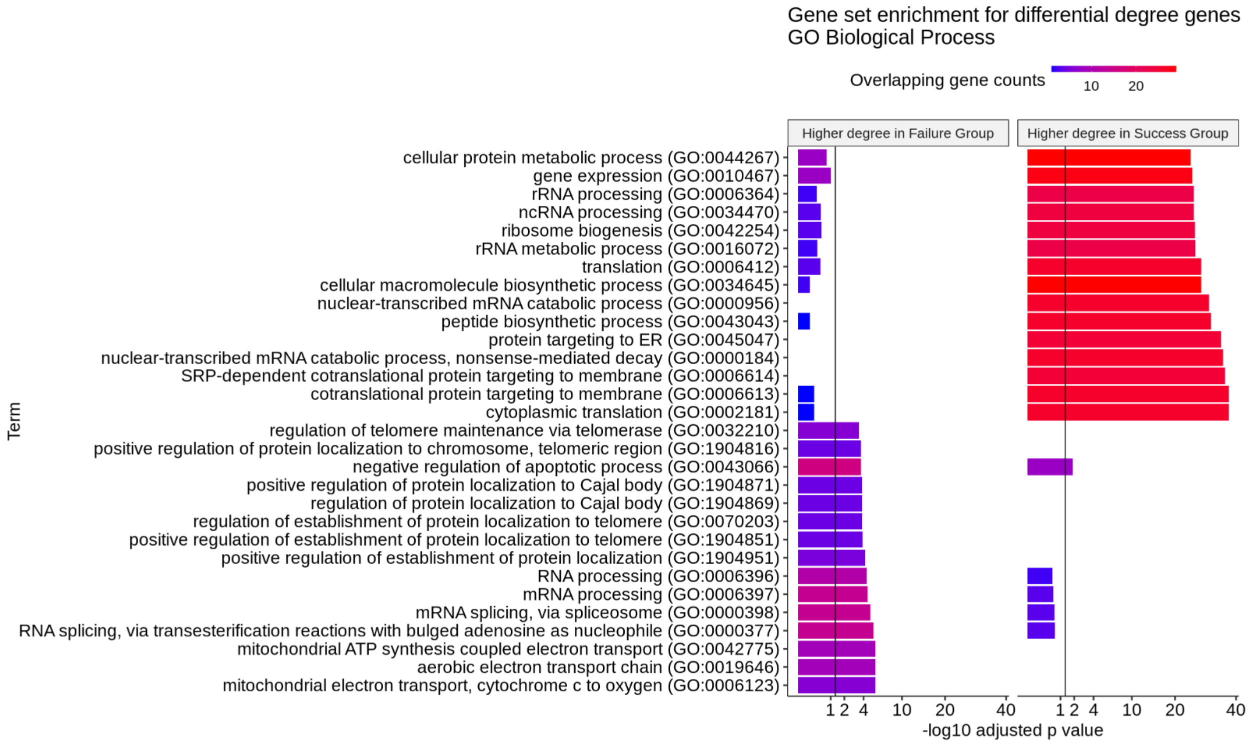


Figure S15: Gene set enrichment analysis of the Dozer identified differential degree genes with GO Biological Processes.

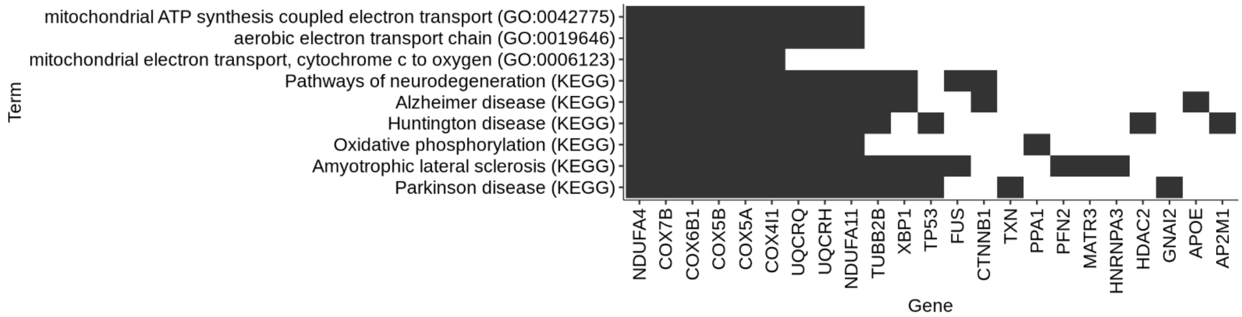


Figure S16: Membership of differential degree genes in KEGG and GO Biological Processes terms (Supplementary Figures S14 and S14). Dark shade indicates that the gene is a member of the relevant KEGG or GO Biological Processes gene set.

### 3 Additional results for the analysis of the `Morabito_2021` dataset

To validate the personalized co-expression networks obtained for the AD and Control donors with data from independent cohorts, we carried out a largest clique analysis. A clique (fully connected sub-network) in the co-expression network represents a group of genes with high co-expression. We computed the largest clique (largest fully connected sub-network) for each donor's network and assessed the consistency of this network structure among donors. The number of genes in largest cliques ranges from 9 to 42. Cliques from different donors have a markedly large proportion of overlapping genes for methods Dozer, Saver, SCT.Pearson and SCT.Spearman, with a mean Jaccard Index of 0.27 to 0.31 (Supplementary Figure S18). Next, for each method, we generated a *clique-set* by pooling genes present in at least two donor largest-cliques. Clique-sets from Dozer, Saver, SCT.Pearson, and SCT.Spearman have higher proportion of gene pairs validated by the STRING PPI database compared to other methods, with an average validation proportion of 0.12-0.17 (Supplementary Figure S18), compared to a baseline of 0.027 among all gene pairs in the network. Clique-set from Dozer network also has high connectivity in an independent single cell RNA-seq of oligodendrocytes from [1], where the average number of edges in the Dozer clique-set is 12 times (22 times) larger than the average connectivity of the whole network for control (diagnosis) donors. Overall, the analysis of largest cliques in donor-specific networks supports that Dozer, Saver, SCT.Pearson, and SCT.Spearman can consistently identify highly co-expressed gene sets in line with protein-protein interactions annotated in the STRING PPI database.

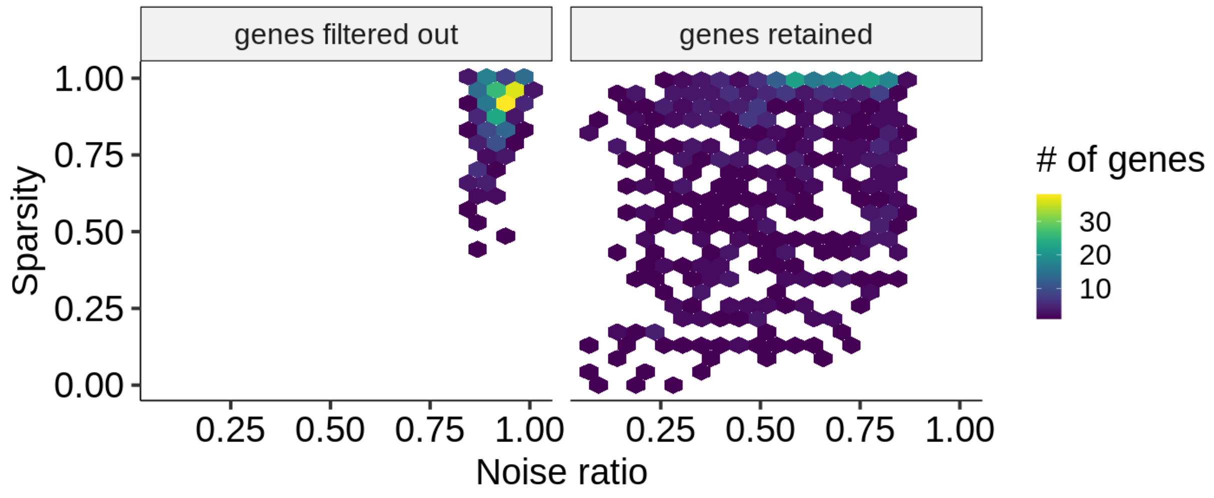


Figure S17: In the Morabito_2021 dataset, 682 out of 1,252 genes were filtered out due to high sparsity and/or high noise ratio. The hexbin plots display the distribution of sparsity levels (proportion of zero counts) and noise ratios among the genes filtered out (left) and the genes retained (right).

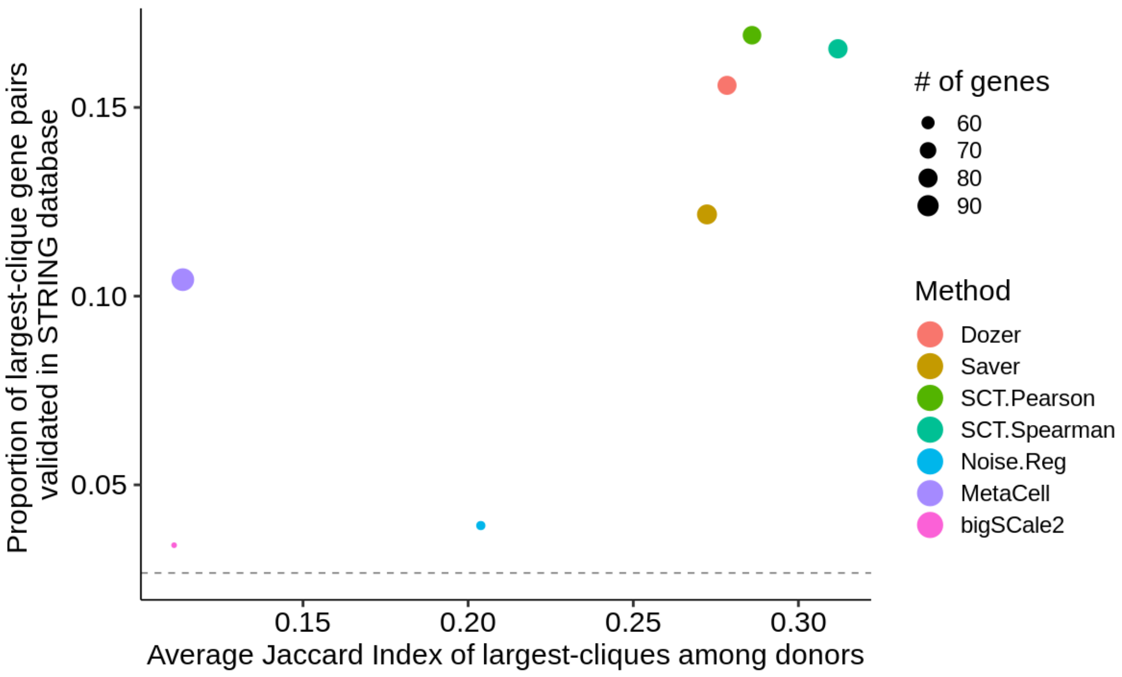


Figure S18: A largest-clique is computed from each donor for each network construction method. The reproducibility of largest-cliques among donors is evaluated using the Jaccard Index (JI) between each pair of donors. X-axis depicts the average Jaccard Index among all donor pairs. The largest-cliques were also evaluated using the STRING database by generating a clique-set across donors for each method. Y-axis depicts the proportion of gene pairs in the clique-set and are validated by the STRING database. The horizontal dashed line at 0.027 represents the STRING database validation rate for a random set of gene-pairs. The size of data points represents the number of genes in the clique-sets, ranging from 51 to 98.

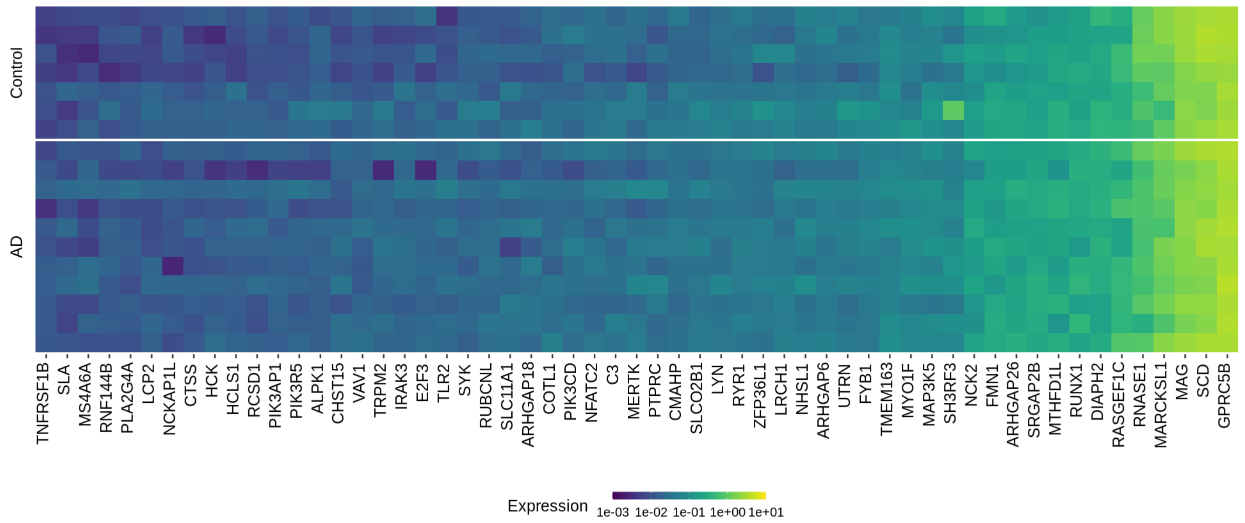


Figure S19: Heatmap of the average expression (normalized expression via SCTransform) of genes in module Dozer-A3 over cells in each donor. Rows and columns in the heatmap correspond to donors and genes, respectively.

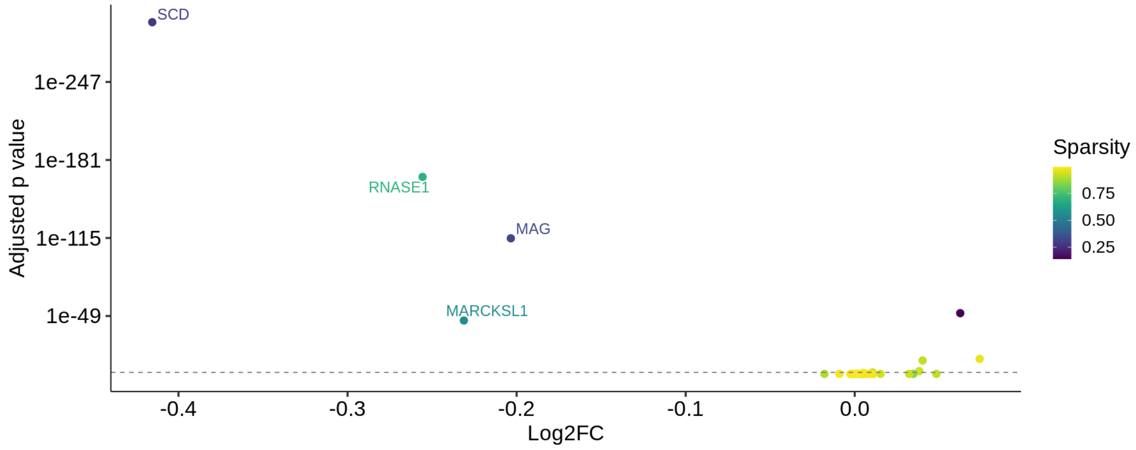


Figure S20: Volcano plot from the differential expression analysis of genes in module Dozer-A3. X-axis represents the log2 fold change of expression in AD versus Control. Y-axis represents adjusted p-value from the differential expression test, where the threshold of 0.05 is marked with the dashed line. Sparsity of the genes is also indicated by color of data points.

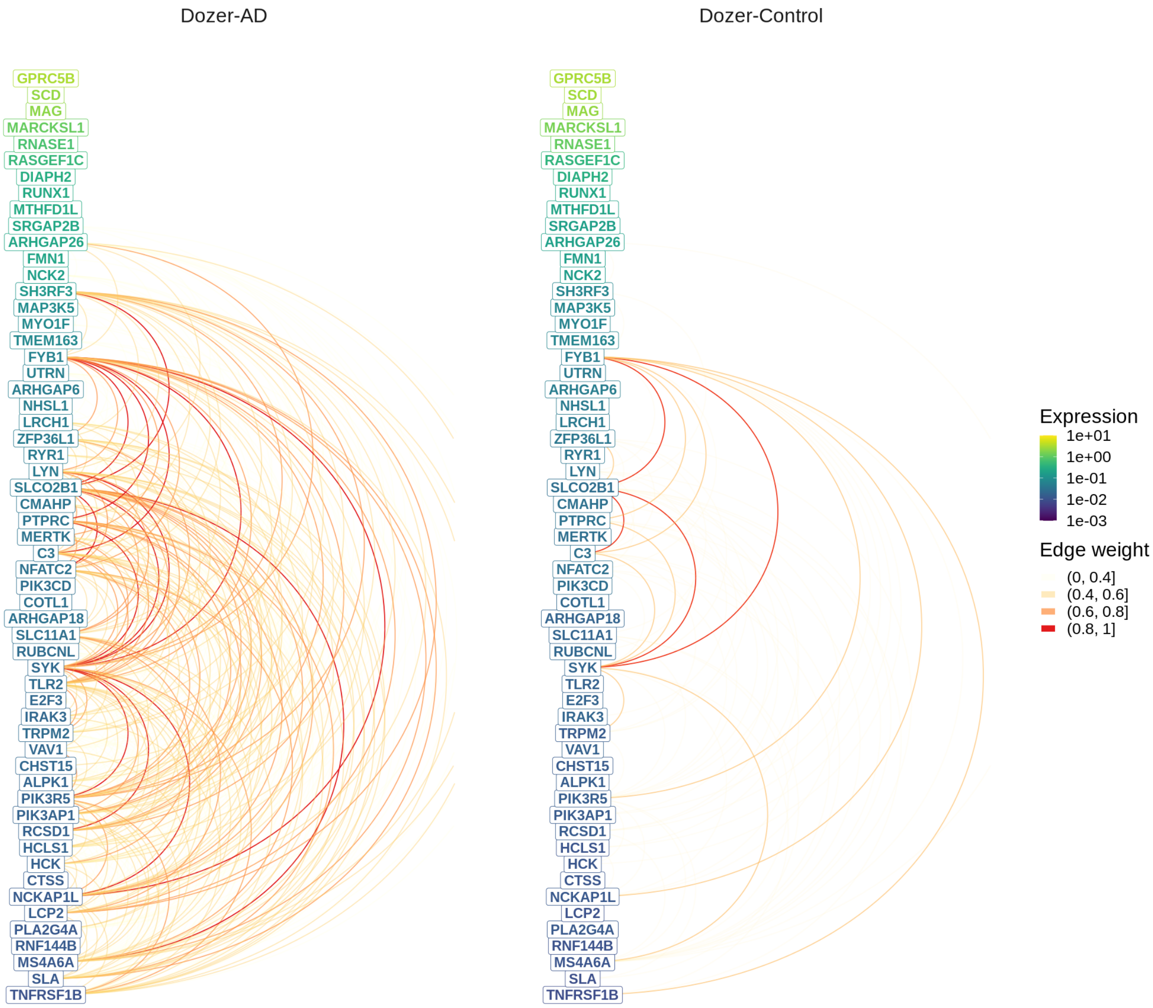


Figure S21: Hive plot visualization of module Dozer-A3 in AD and Control groups. Genes are ordered from high (top) to low (bottom) by their average expression across all donors, and colored by the average expression in the corresponding diagnosis group. The arcs between the genes in this linear layout depict the edges in the average Dozer networks of AD (left) and Control (right) donors, with colors representing the edge weights.

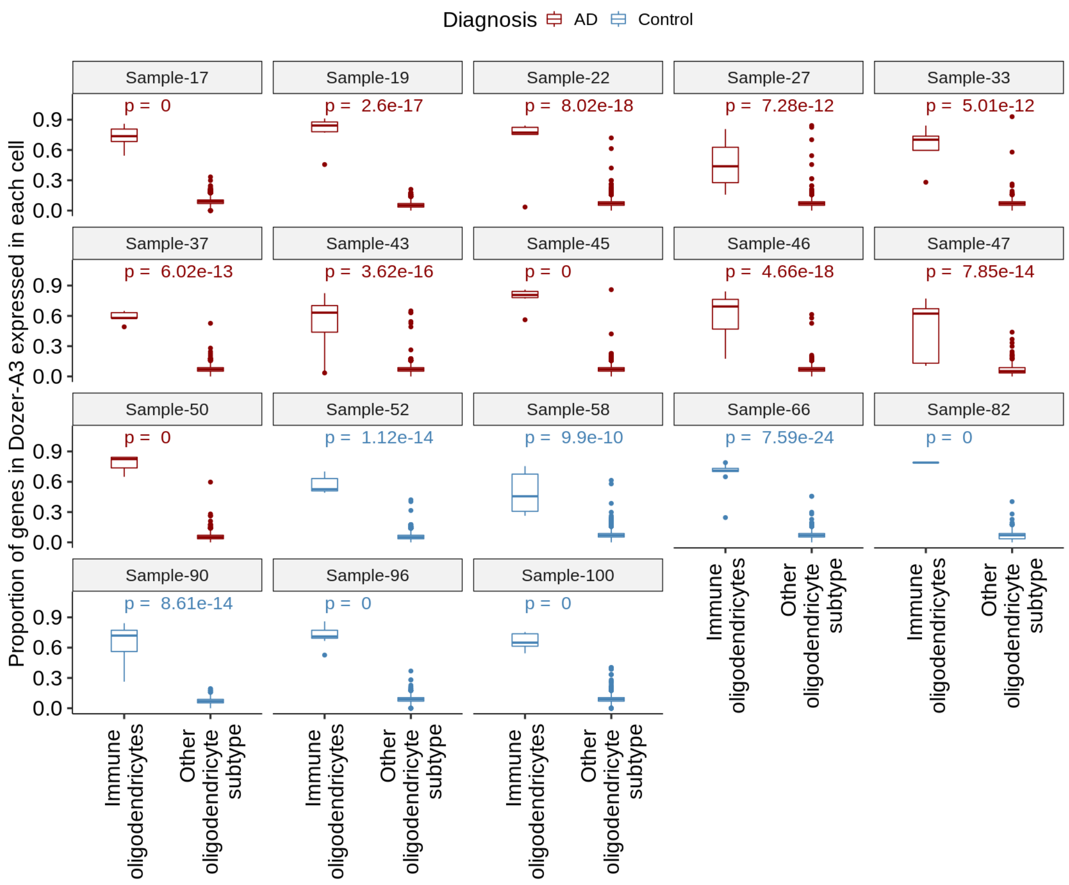


Figure S22: Proportion of the Dozer-A3 genes expressed in each cell at the donor level. Individual cells in each of the two cell populations, immune oligodendrocytes (ODC) and other ODC subtypes, are evaluated with respect to the proportion of expressed Dozer-A3 genes at the donor level. The reported p-values for each donor evaluate whether the co-expression of module Dozer-A3 in donor immune oligodendrocytes is higher than expected by chance (Method).

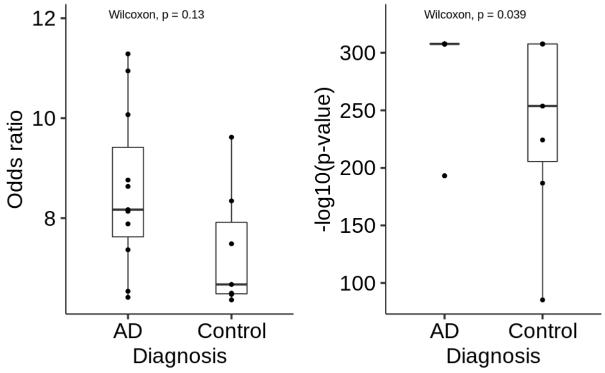


Figure S23: Boxplots of the odds ratios and -log10(p-values) from Fisher's exact tests that evaluate the association between co-expressing Dozer-A3 module genes and being an immune ODC cell for each donor (Method). Wilcoxon rank sum tests were conducted to evaluate the differences in odds ratios and -log10(p-value)s between the two diagnosis groups (AD vs. Control).

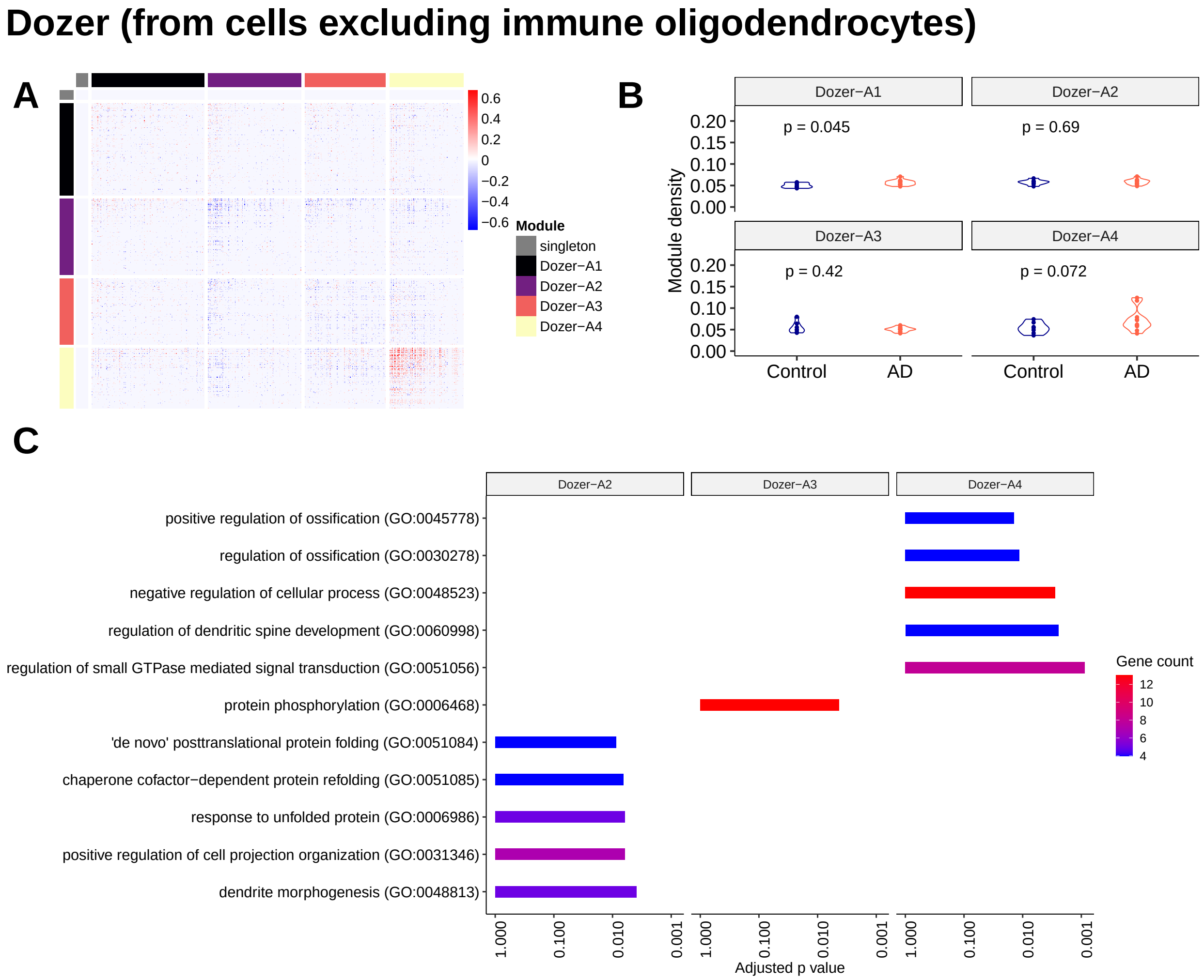


Figure S24: **(A)** Heatmap of gene modules from the Dozer "difference network" constructed with cells excluding immune oligodendrocytes. **(B)** Violin plot of module densities. **(C)** Top 5 significant GO terms from gene set enrichment analysis of each module (modules with no significant terms are excluded).

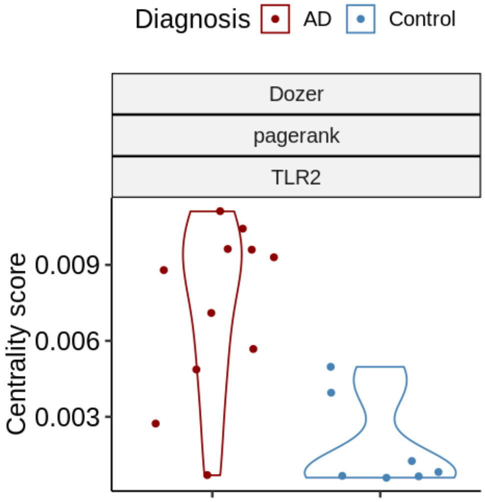


Figure S25: Violin plots of the gene centrality measures in the AD and Control groups for the genes with significant differential centrality at 5% FDR in the `Morabito_2021` dataset. The three grid labels indicate the co-expression network construction method, centrality, and gene name.

### Saver

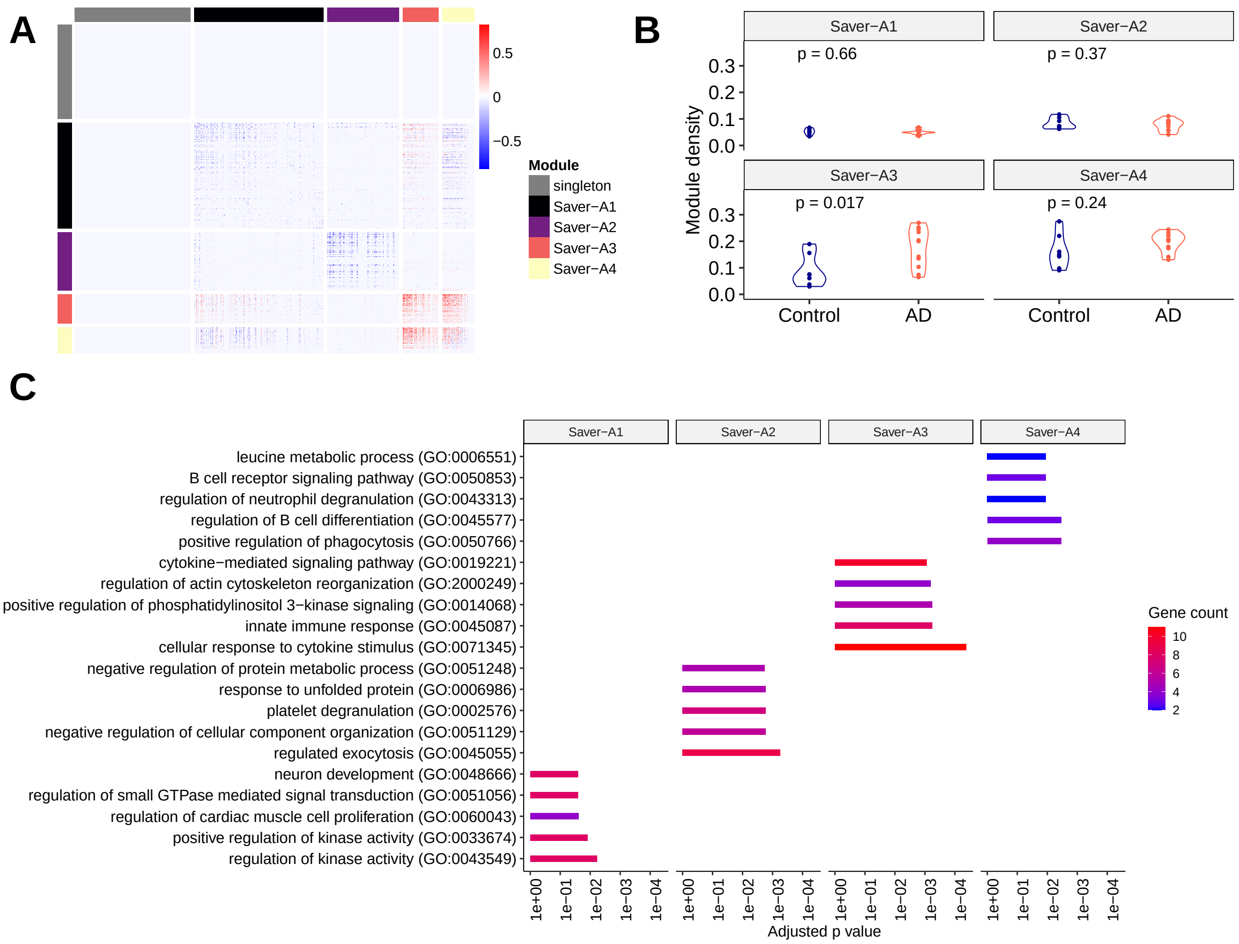


Figure S26: **(A)** Heatmap of gene modules from the Saver "difference network". **(B)** Violin plot of module densities. **(C)** Top 5 significant GO terms from gene set enrichment analysis of each module (modules with no significant terms are excluded).

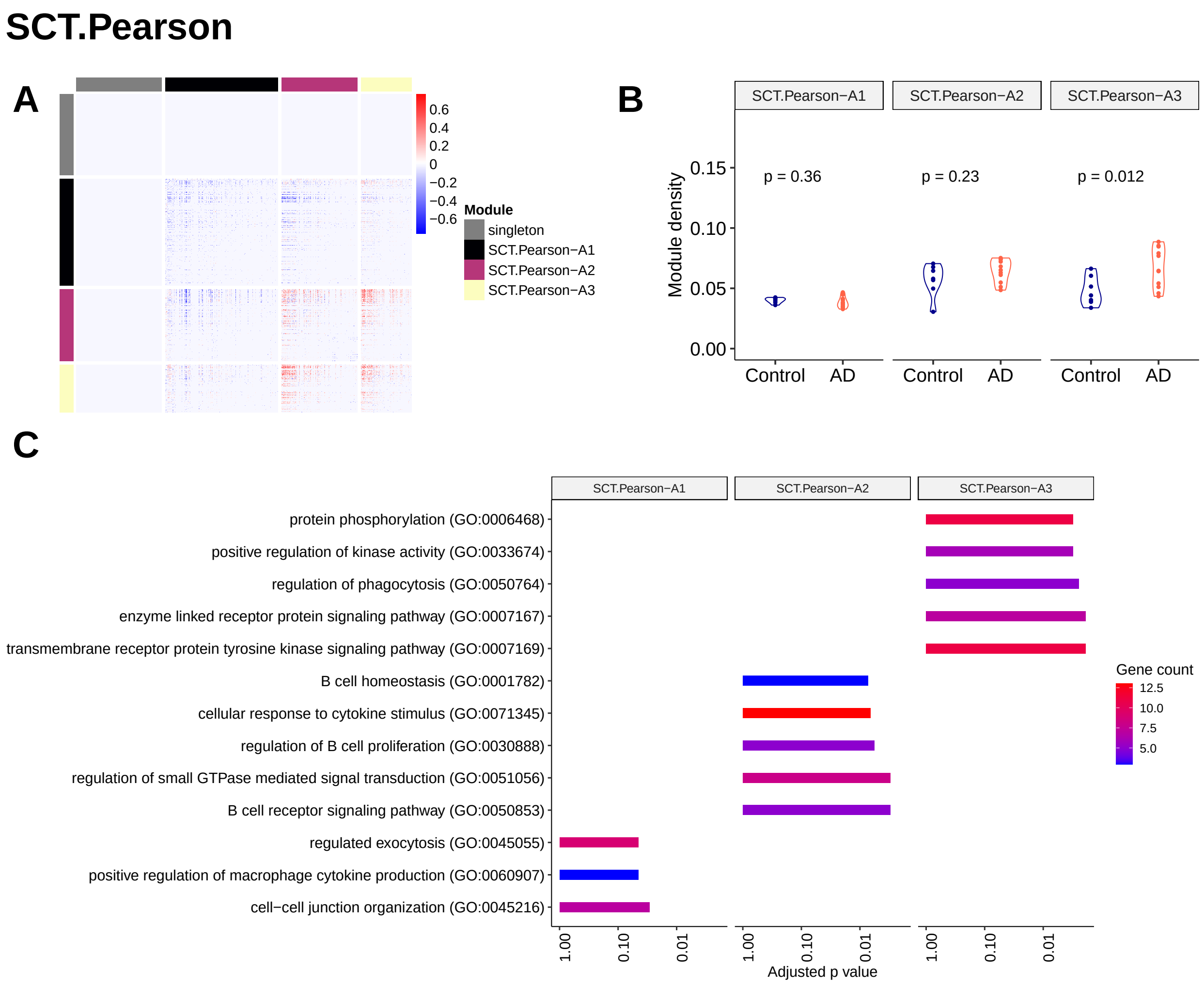


Figure S27: **(A)** Heatmap of gene modules from the SCT.Pearson "difference network". **(B)** Violin plot of module densities. **(C)** Top 5 significant GO terms from gene set enrichment analysis of each module (modules with no significant terms are excluded).

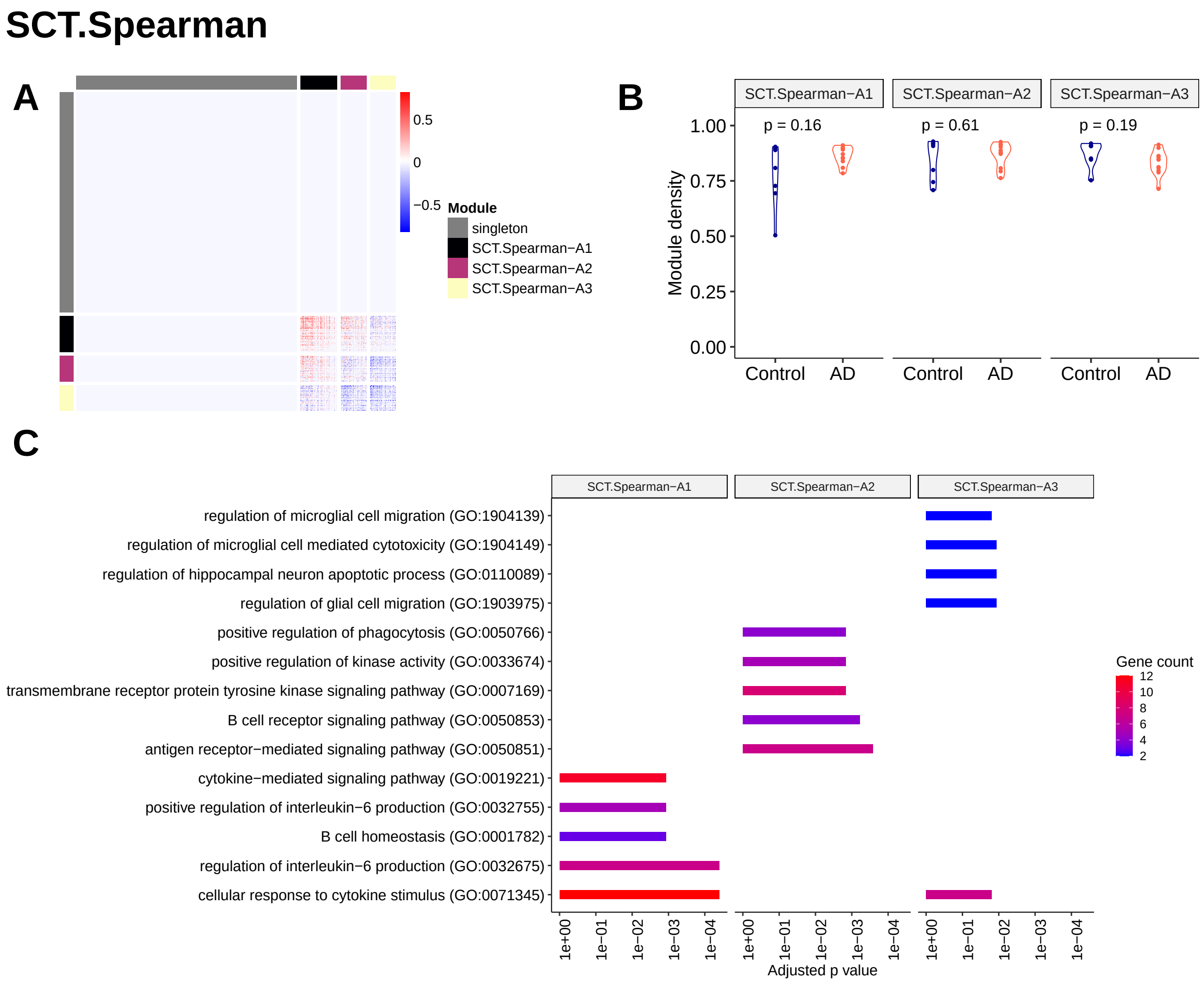


Figure S28: **(A)** Heatmap of gene modules from the SCT.Spearman "difference network". **(B)** Violin plot of module densities. **(C)** Top 5 significant GO terms from gene set enrichment analysis of each module (modules with no significant terms are excluded).

#### Noise.Reg

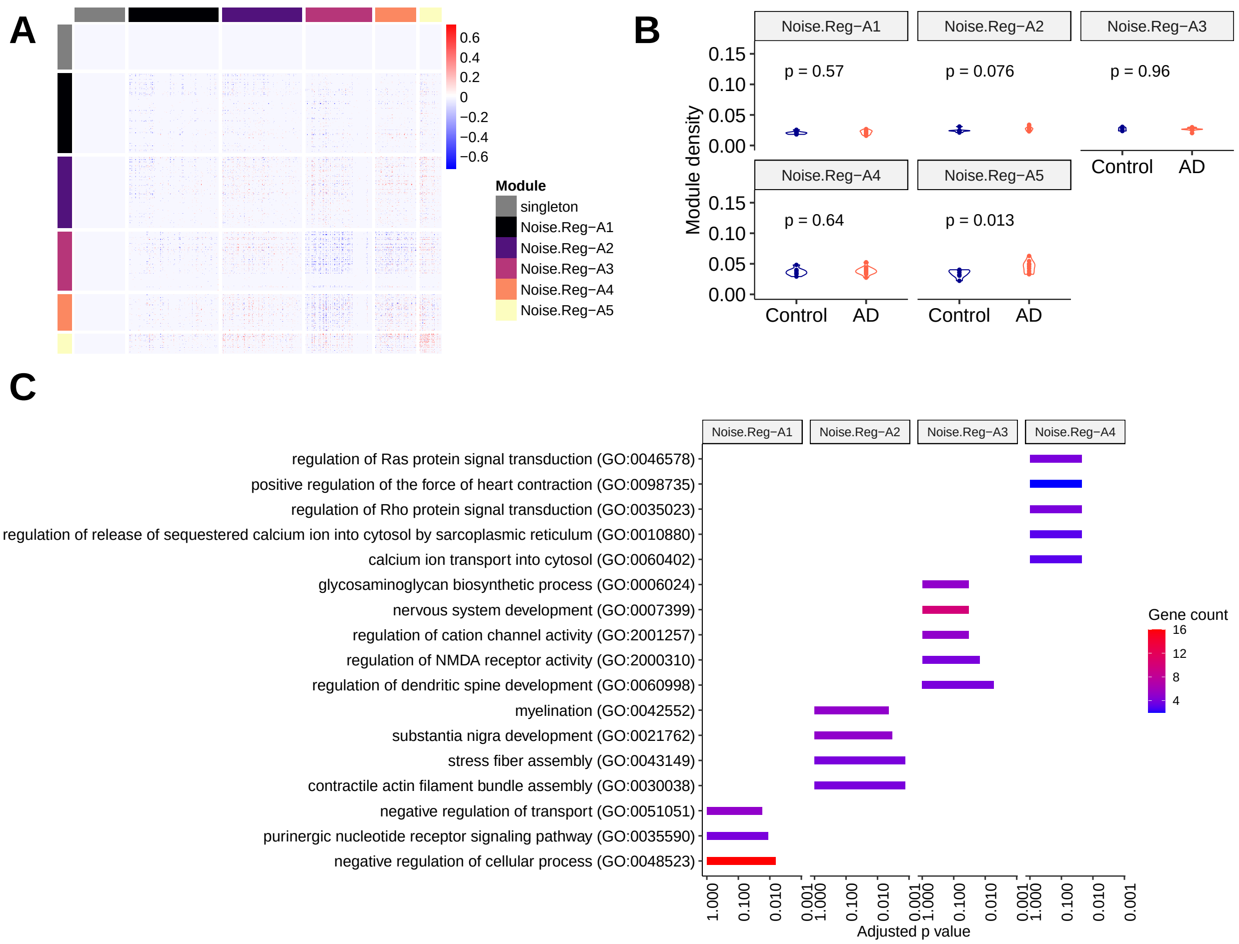


Figure S29: **(A)** Heatmap of gene modules from the Noise.Reg "difference network". **(B)** Violin plot of module densities. **(C)** Top 5 significant GO terms from gene set enrichment analysis of each module (modules with no significant terms are excluded).

#### MetaCell

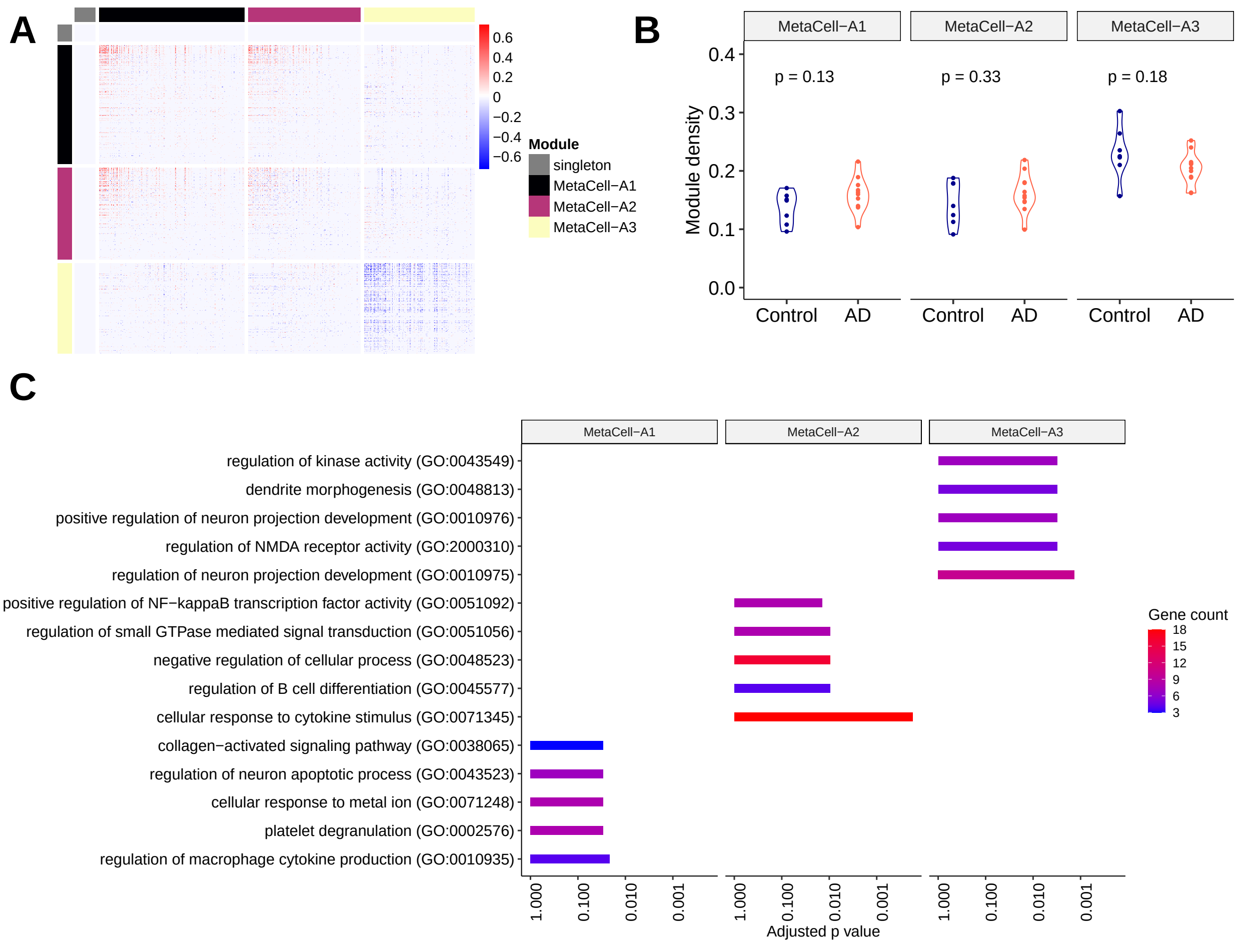


Figure S30: **(A)** Heatmap of gene modules from the MetaCell "difference network". **(B)** Violin plot of module densities. **(C)** Top 5 significant GO terms from gene set enrichment analysis of each module (modules with no significant terms are excluded).

#### bigSCale2

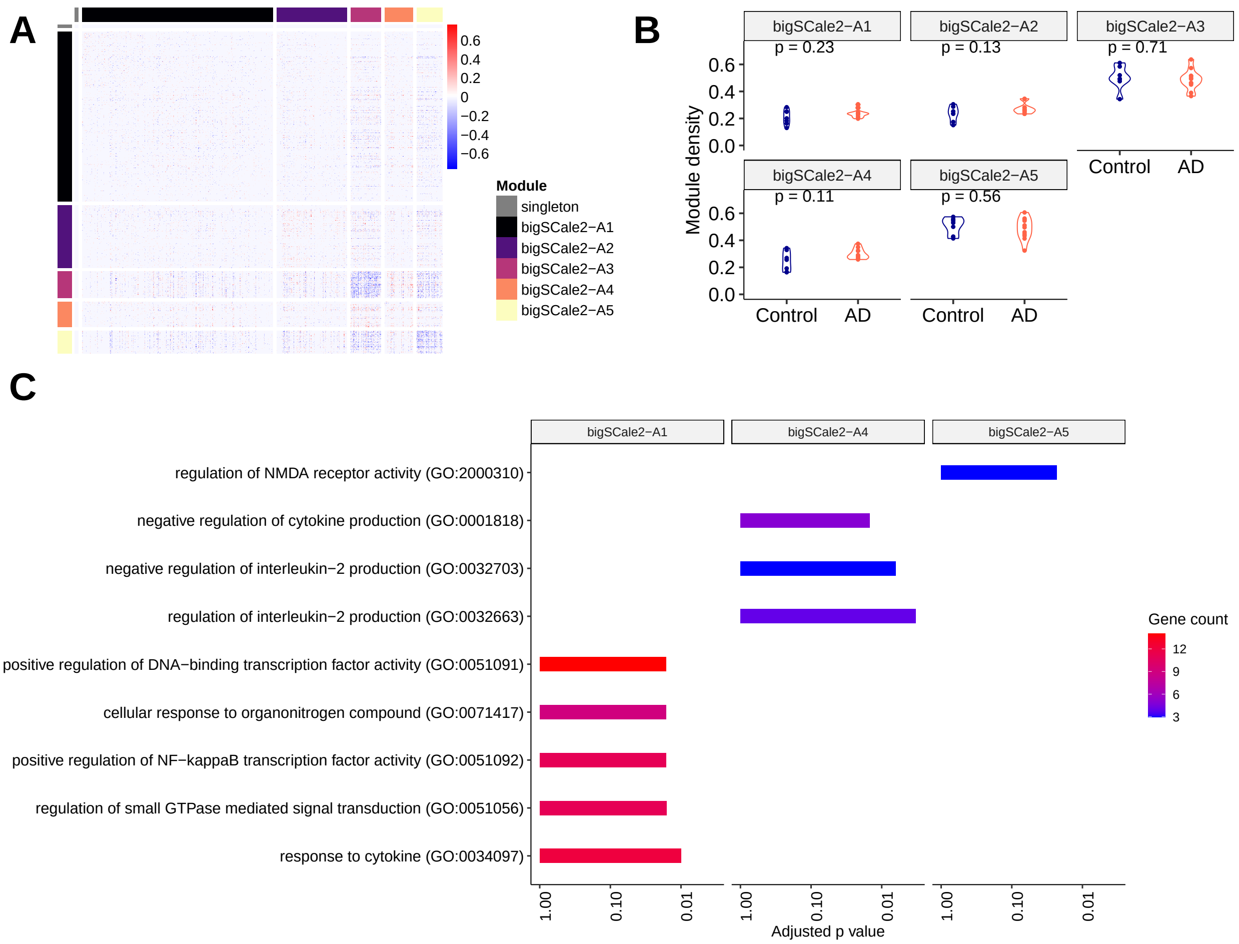


Figure S31: **(A)** Heatmap of gene modules from the bigSCale2 "difference network". **(B)** Violin plot of module densities. **(C)** Top 5 significant GO terms from gene set enrichment analysis of each module (modules with no significant terms are excluded).

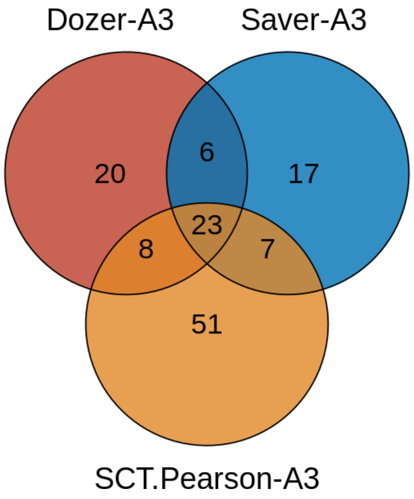


Figure S32: Venn diagram of gene modules identified by Dozer, Saver, and SCT.Pearson and with module densities significantly associated with the AD diagnosis.

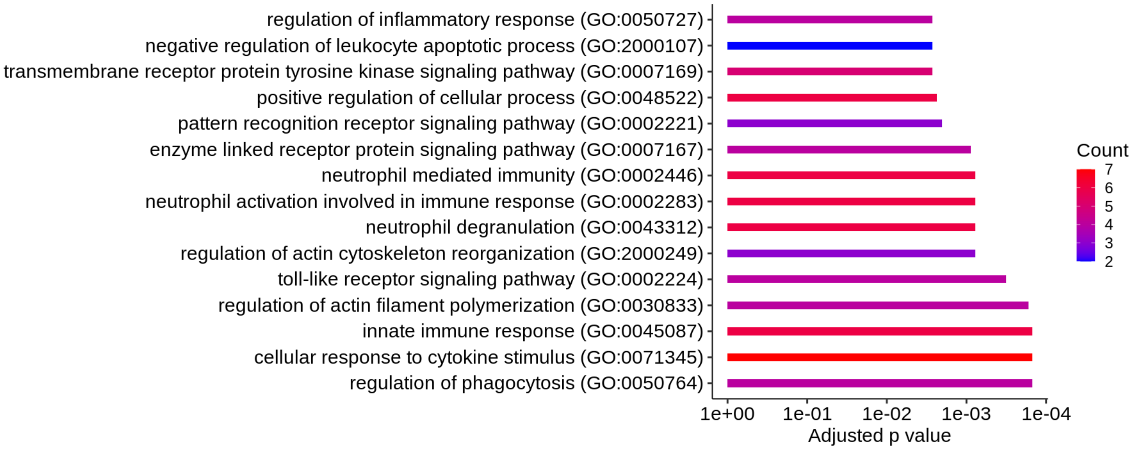


Figure S33: GO enrichment analysis of the genes shared between co-expression modules "Dozer-A3", "Saver-A3", and "SCT.Pearson-A3".

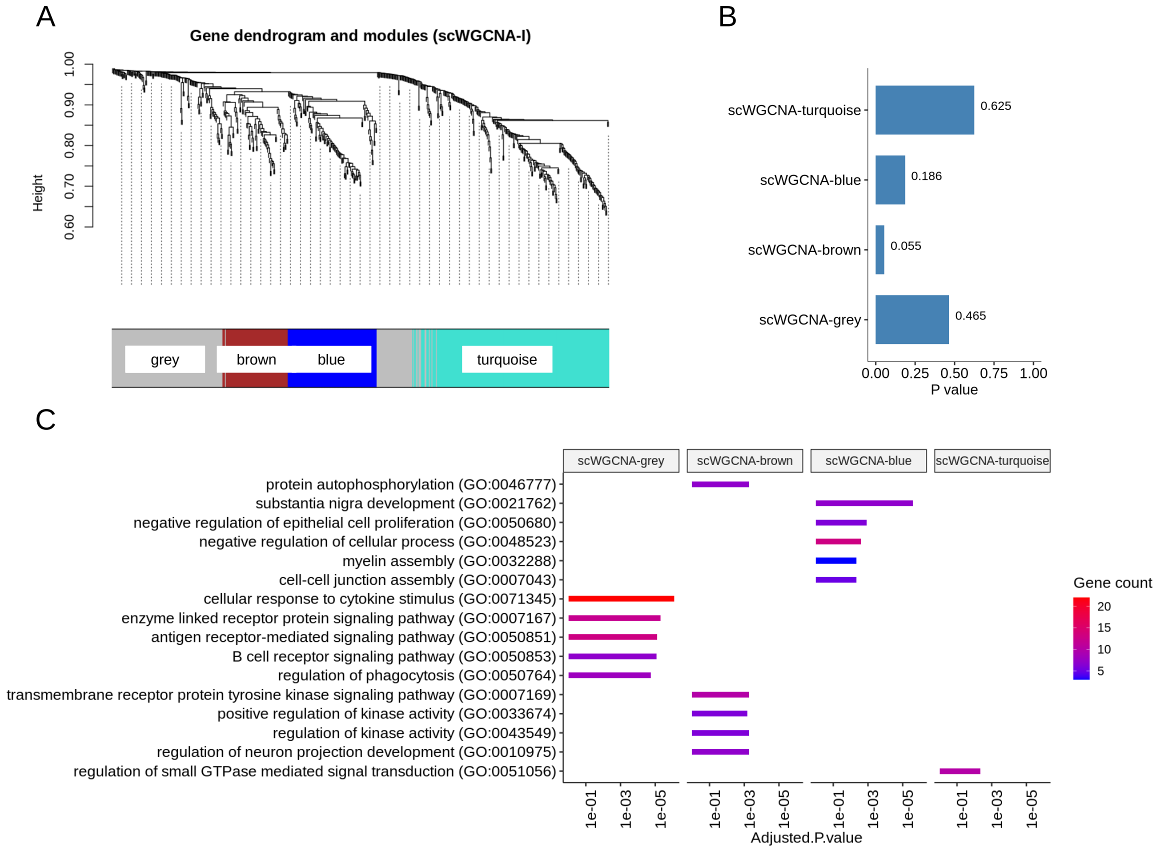


Figure S34: **(A)** Module dendrograms of scWGCNA-I. **(B)** P-values of the association tests between module eigengene expression and diagnosis. **(C)** Top 5 significant GO terms for each gene module.

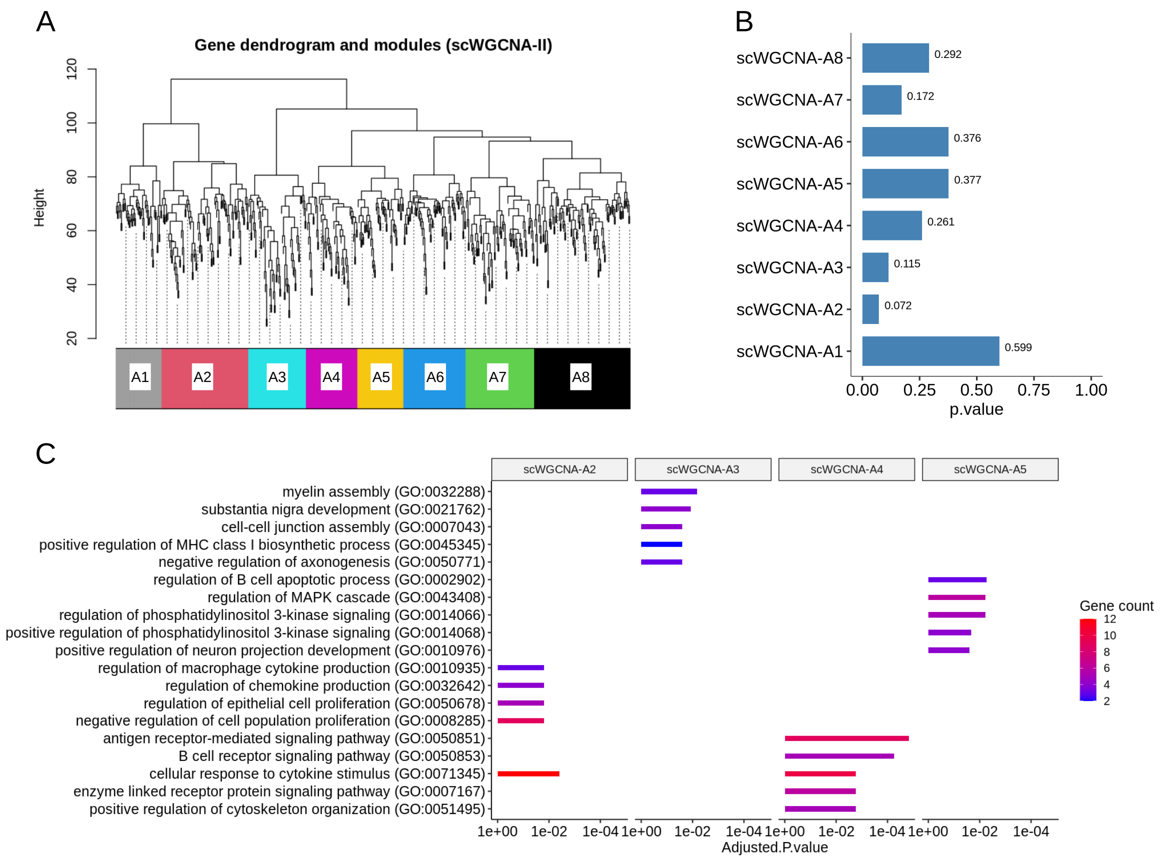


Figure S35: **(A)** Module dendrograms of scWGCNA-II. **(B)** P-values of the association tests between module eigengene expression and diagnosis. **(C)** Top 5 significant GO terms each gene module (modules with no significant terms are excluded).

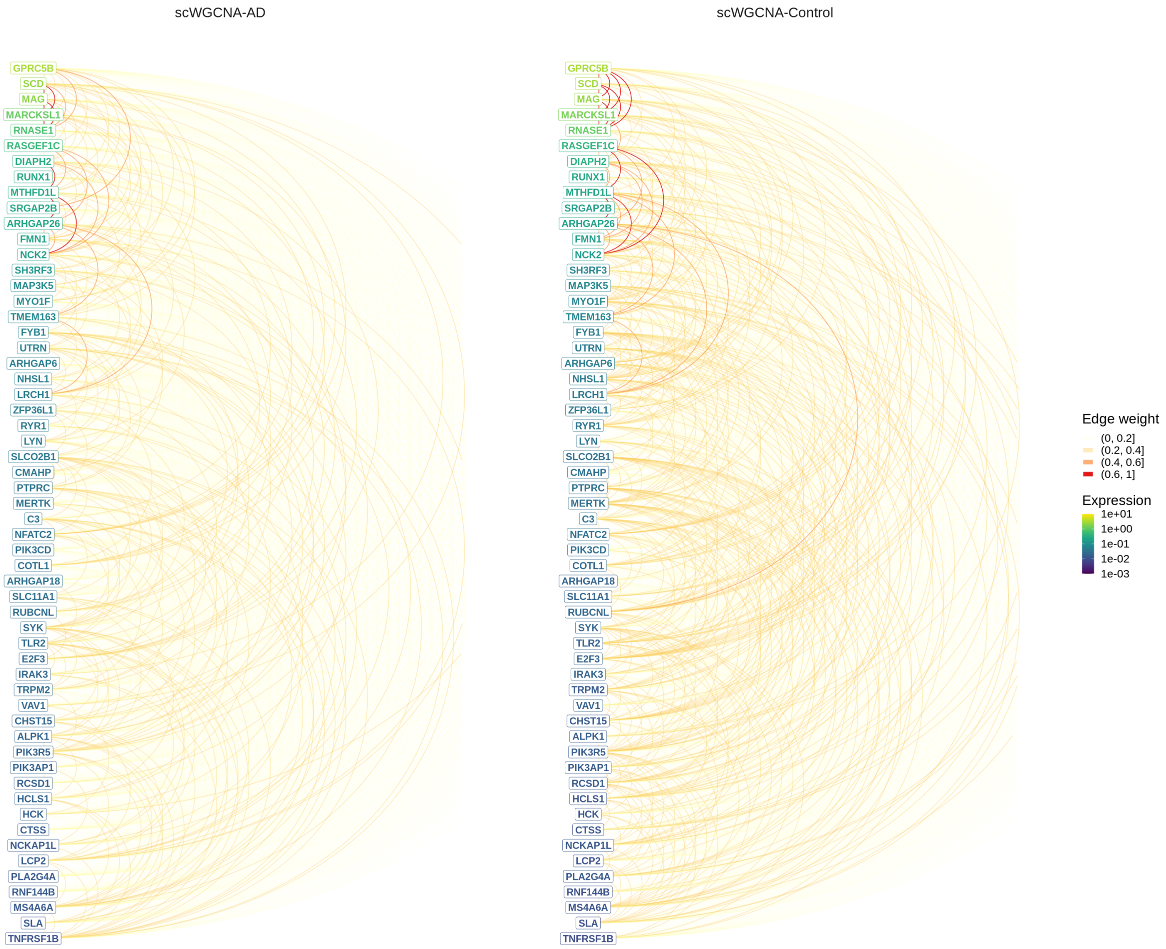


Figure S36: Hive plot visualization of scWGCNA gene correlation matrices for module Dozer-A3 in Control and AD groups. Genes are ordered from high (top) to low (bottom) by their average expression across all donors, and colored by the average expression in the corresponding diagnosis group. The color of arcs between the genes in this linear layout depicts the absolute correlation between genes in scWGCNA networks of AD (left) and Control (right) donors.

### References

[1] Nagy C, Maitra M, Tanti A, Suderman M, Théroux JF, Davoli MA, Perlman K, Yerko V, Wang YC, Tripathy SJ, et al.. 2020. Single-nucleus transcriptomics of the prefrontal cortex in major depressive disorder implicates oligodendrocyte precursor cells and excitatory neurons. *Nature neuroscience* **23**: 771–781.

[2] Sun T, Song D, Li WV, and Li JJ. 2021. scDesign2: a transparent simulator that generates high-fidelity single-cell gene expression count data with gene correlations captured. *Genome biology* **22**: 163.

[3] Zappia L, Phipson B, and Oshlack A. 2017. Splatter: simulation of single-cell RNA sequencing data. *Genome biology* **18**: 174.
